# Enhancer-driven gene regulatory network of forebrain human development provides insights into autism

**DOI:** 10.1101/2023.09.06.555206

**Authors:** Alexandre Jourdon, Jessica Mariani, Abhiram Natu, Feinan Wu, Boxun Li, Davide Capauto, Kevin T. Hagy, Scott Norton, Livia Tomasini, Alexias Safi, Anahita Amiri, Jeremy Schreiner, Cindy Khanh Nguyen, Neal Nolan, Matthew P. Nelson, Daniel M. Ramos, Michael E. Ward, Anna Szekely, James C. McPartland, Kevin Pelphrey, Pamela Ventola, Katarzyna Chawarska, Charlie A Gersbach, Gregory E. Crawford, Alexej Abyzov, Flora M. Vaccarino

## Abstract

Cell differentiation is orchestrated by transcription factors (TFs) binding to enhancers, shaping gene regulatory networks that drive neuronal lineage specification. Deciphering these enhancer-driven networks in human forebrain development is essential for understanding the genetic basis of neurodevelopmental disorders. Through integrative epigenomic and transcriptomic analyses of human forebrain organoids derived from 10 individuals with autism spectrum disorder (ASD) and their neurotypical fathers, we constructed a comprehensive enhancer-driven gene regulatory network (GRN) of early neurodevelopment. This GRN revealed hierarchical regulatory transitions guiding neuronal differentiation and was experimentally validated via CRISPR interference (CRISPRi) and loss-of-function analyses. A subnetwork linked ASD-associated transcriptomic alterations to dysregulated TF activity, implicating FOXG1, BHLHE22, EOMES, and NEUROD2 as key regulators of excitatory neuron specification in macrocephalic ASD. These findings suggest that ASD disrupts enhancer-driven regulatory frameworks, altering neuronal cell fate decisions in the developing fetal brain.

## Introduction

Fundamental cell fate decisions and regional specification during early human brain development underlie our larger brain size, cellular diversity, and cognitive abilities, but also our susceptibility to neurodevelopmental disorders, such as autism spectrum disorders (ASD). Brain organoids derived from human induced pluripotent stem cells (iPSC) are a multilineage model that emulates the dynamics of brain development in an individual’s genetic background^1–3^. While several transcriptomic studies have charted lineage specific gene expression patterns of fetal human brain and brain organoids, less is known about how those lineages are specified by the non-coding genome, and specifically by enhancers, the main elements implicated in the modulation of gene expression^4^. Located at variable distances from gene promoters, enhancers are bound by specific transcription factors (TFs) which can modify chromatin conformation and interact with the promoter machinery to increase or decrease the expression of downstream genes ^5–7^. TFs, enhancers and other types of regulatory regions have been described in the adult^8,9^ and prenatal ^10–12^ human brain, but postmortem brain studies cannot capture the dynamic unfolding of developmental events and do not allow for experimental manipulation. We and other have started to chart enhancers location and activity during early brain development in both fetal human cortex and isogenic organoids ^13^ and a recent study has used the organoid approach to identify the regulatory events that drive early cell fate decisions between neural and non-neural cells ^14^. Understanding the regulatory relationships between enhancers, TFs and gene expression during development remains an important step to understand the etiology of neurodevelopmental disorders.

In our recent study of cortical organoids from 13 families with idiopathic ASD with or without macrocephaly, we found by scRNA-seq analyses a strong imbalance in excitatory neuron lineages when comparing ASD probands to their neurotypical father^15^. In the present study we performed histone marks chromatin immunoprecipitation and sequencing (ChIP-seq) and parallel RNA sequencing analyses in iPSC-derived forebrain organoids from 10 out of those 13 families. Integrating enhancer activity with TF binding sites and gene expression on a genome-wide scale, we built a gene regulatory network (GRN), identifying enhancers and cognate TFs active during the progression of neurogenesis and the diversification of neuronal subtypes. By comparing enhancer activity and differential gene expression of individuals with ASD to their unaffected father as controls, we further delineated the regulome underlying the transcriptional alterations observed in ASD with and without macrocephaly.

## Results

### 1. Building a gene regulatory network based on epigenome and transcriptome during forebrain organoid development

#### Study cohort

In the present study we characterize epigenomic and transcriptomic states of precursor cells and early neurons using iPSC lines from 10 idiopathic ASD individuals and their unaffected fathers differentiated into forebrain organoids and analyzed by scRNA-seq, ChIP-seq and bulk RNA-seq. The clinical information and genetic background of the individuals have been previously reported in a single cell transcriptome study of this cohort ^15^; we now analyze ChIP-seq and bulk RNA-seq data collected in parallel to understand the regulatory events underlying the transcriptomic phenotypes observed in the ASD individuals. ASD probands were classified as “macrocephalic” if exhibiting head size above 90^th^ head size percentile or as “normocephalic” if exhibiting normal head size using a normative dataset^16^. Since macrocephaly is often a family trait, the control fathers of macrocephalic probands were also macrocephalic (see clinical information in **Table S1**). The iPSC lines from all individuals were previously characterized by whole genome sequencing^15^; although some probands carried a variant affecting a gene associated with syndromic ASD, except for a large duplication, variants did not change the expression for the corresponding genes, and variants did not converge on the same genes across our idiopathic cohort (see Jourdon et al, 2023 for full description of the results).

All iPSC lines were differentiated into forebrain organoids and histone marks-ChIP-seq (H3K4me3, H3K27ac, H3K27me3 and H3K4me1), bulk RNA-seq and scRNA-seq were concurrently performed at terminal differentiation day 0 or TD0 (corresponding to 20 days after neural induction of the iPSC lines), TD30 and TD60 (**Fig. 1A, Table S1**). We then developed a pipeline that integrated this multi-omics data collected in parallel into a *regulatory network* of human forebrain organoid development (**Fig. 1B, Methods**). First, integrated annotation of the genome using a ChromHMM 9-states model^17^ for ChIP-seq data identified putative promoters, enhancers (∼360,000), and silent chromatin regions (**Fig. 1C-D).** Enhancers were then linked to potential target genes by proximity to promoters (20 kb) or by reported distal interactions identified in human fetal brain Hi-C datasets^18–20^ **(Fig. S1A).** Altogether, 173,437 gene-linked enhancers (GLEs) were mapped across all samples (**Fig. S1B-E, Table S2**).

**Figure 1:**
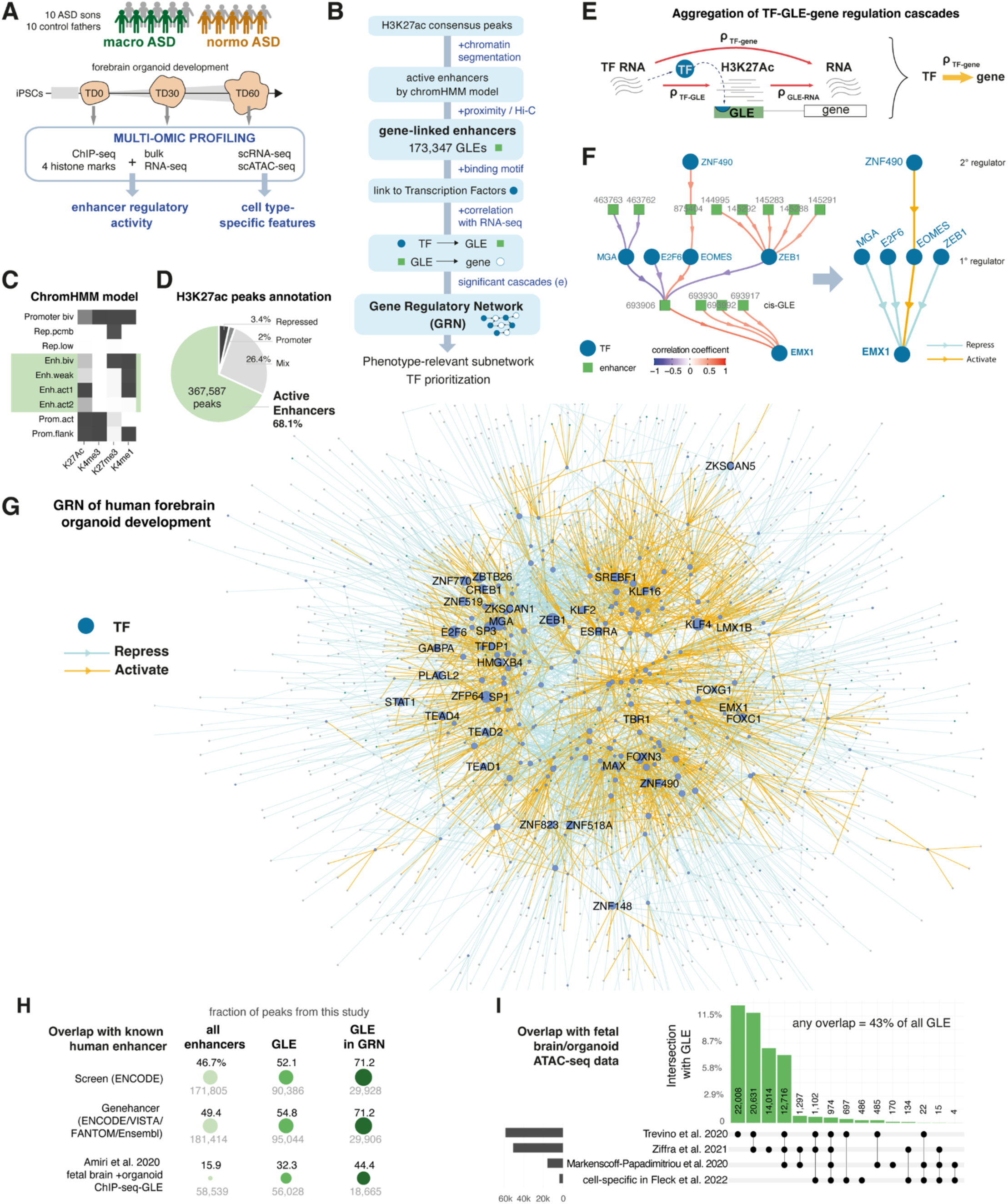
Building a regulatory network during forebrain organoid development anchored around the identification of active enhancer elements. **A.** Study design with multi-Omic characterization of forebrain organoid development in 20 individuals including ASD individuals with or without macrocephaly (“macroASD”, head circumference > 90^th^ percentile of the population, “normo ASD” otherwise). **B.** Data integration pipeline to generate the GRN (see **Methods**). It incorporates enhancer-gene linkages from proximity and Hi-C, TFBS identification, and correlation of enhancer activity with expression of upstream TF(s) and downstream gene(s). The final step consists in the derivation of a GRN through aggregation of the TF-enhancer-gene cascades (see **E**). **C.** The 9-states model of genome annotation built using the 4 chromatin marks ChIP-seq data from this study. Four enhancer states were then merged into an “active enhancer” state. Enhancers profiles were essentially characterized by H3K27Ac without H3K4Me3 activity. **D.** Diagram showing the proportion of each main chromatin state identified in (**C**) for all H3K27ac consensus peaks. **E.** Schematic of the aggregation strategy validating the intermediate role of enhancers through concordant correlation at the RNA level between TF and downstream genes (see also **Fig. S2**). Regulatory actions were then summarized to build the TF-gene GRN, conserving only the correlation between TF and gene in the final network. Blue arrows=negative and red arrows= positive correlation (*ρ*). **F.** Upstream enhancer regulatory network of the gene EMX1. Arrows indicate regulatory link between TFs and their bound GLEs (in green with unique ID from **Data S2**) and links from GLE to their downstream gene in *cis*. Colors of the arrows show correlation level between the linked elements. Note the four-steps distance between the TF ZNF490 and EMX1 expression. The corresponding TF-gene network is plotted on the right using TF-gene RNA level correlation as link weight (two-sided Spearman’s correlation FDR corrected p-value < 0.05). **G.** Visualization of the central component of the GRN inferred from the enhancer regulatory network, plotted by Kamada-Kawai graph layout. Genes connected by repressive regulation (i.e., *ρ*_*TF–RNA*_ < 0) are further apart than genes connected by activating regulation (see **Methods**). By construction, TF (blue dots) are central element of the GRN, each connecting in their regulon 1 to 1,047 downstream genes (small grey dots). Top 50 TF by regulon size (dot size) and salient TFs from our study are named. **H** Fraction of all enhancers regions (“All Enh.”), gene-linked enhancer (“GLE”) and regulatory GLE (“reg-GLE”, i.e., GLE significantly correlated in activity with gene expression, FDR < 0.01, **Data S2**) that overlap with reported human enhancers from 3 sources ^13,22,23^. Dot shows percentage enhancer from our study overlapping with each source, corresponding number of peaks is indicated below. All overlaps increased significantly from “All Enh.” to “GLE” to “reg-GLE” (p-values of one-tailed Fisher exact test < 2.2 x 10^-16^). **I.** Upset plot showing the intersection between GLE from this study and peaks identified in published bulk and single-cell ATAC-seq data from human fetal brain or brain organoid models ^10,14,24,25^.

GLEs were then linked to TFs by the presence of a TF binding site (TFBS) motif in their sequence (**Fig. S1E-F)**. Next, significant correlations between GLE activity (H3K27ac reads) and expression of the linked gene (RNA reads) across samples were used to retain the most functionally relevant links and infer the direction of the regulation. Both activating (positive correlation) and repressing (negative correlations) regulatory relationships linking TFs to enhancers, and enhancers to their downstream genes were retained (**Fig. 1B, Fig. S2A,B**). Repressive GLE-genes regulatory relationships (n=1,556) were much fewer than activating ones (n=6,794). Repressive and activating TF-GLE regulatory relationships were similar in numbers, suggesting that ∼50% of developmental TF could act as repressors. While we could not exclude that part of this negative correlation could be explained by coincidental absence of activators, repressor TFs are known to be able to induce histone deacetylation and inhibit the recruitment of basal machinery without inducing chromatin compaction^21^. We therefore decided to incorporate both positive and negative TF-to-enhancer regulatory links in our network analysis.

All retained links were integrated into a TF-enhancer-gene network connecting 373 TFs, 42,035 GLEs, 6,501 protein coding genes and 1,690 long non-coding RNAs (lncRNAs) (**Table S2, T2-3**). In such network, active enhancers are pivotal elements in the upstream regulatory network of any expressed gene, some with complex regulatory cascades (**Fig. 1E-F, Fig. S3**). Compared to initial set of GLEs called, the subset of GLEs included in the GRN showed a significantly higher overlap with sets of known enhancers in human ^13,22,23^ (71% overlap with ENCODE candidate regulatory elements), confirming the relevance of using correlation with transcription levels to better refine regulatory elements identification (**Fig. 1H)**. Many identified GLEs also overlapped with open chromatin regions identified by bulk and single-cell ATAC-seq assays performed on human fetal brain or brain organoids in published datasets ^10,14,24,25^ (43% overlap, **Fig. 1I**). This degree of overlap was remarkable considering that ChIP-seq and ATAC-seq identify regulatory regions based on histone post-translational modification vs accessible chromatin region.

To capitalize on this characterization of enhancer-centered regulatory cascades, we derived a reduced representation in the form of a gene regulatory network (GRN), where TF were directly connected to downstream gene(s) when one or several intermediate functional GLE(s) linking them was identified (**Fig.1E-F, Fig. S2C**). We pruned 13.3% of such links by requiring a significant correlation not only between TF expression and GLE activity and GLE activity and target gene(s), but also between expression of the TF and downstream gene and used the obtained correlation coefficient to infer both the strength and the activating or repressing nature of the TF-to-gene regulatory action (**Methods)**. The resulting GRN consisted of 330 upstream TFs regulating 5,067 downstream transcripts, encompassing 4,034 protein-coding genes and 1,023 lncRNAs (**Fig. 1G, Table S3**). As we incorporated data from both ASD probands and controls to increase confidence in correlation inference, we verified that ASD diagnosis had a limited effect on the variance of gene and TF’s expression and was unlikely to drive correlation results (**Fig. S2D,E**), ensuring that the GRN was representative of molecular events driving typical neurogenesis. We found a limited overlap between the obtained GRN and a published regulatory network derived by integrating scATAC-seq and scRNA-seq conducted on stem cells, early progenitor cells and organoids ^14^ (2.3% of TF-gene regulatory edges in common). The inclusion of earlier time points and methodological differences suggested that the two networks described connections of different sets of TFs (200 TFs in common or 23.5% of the union) and genes (823 targets or 12% of the union) (**Fig. S4C-F**). However, the large majority of the common edges (239/249) show consistent measures in TF-to-gene correlation estimates between the two studies, suggesting that both networks captured the same regulatory interactions (**Fig. S4F**). Ultimately, those different networks could be used in complement to chart a more complete view of regulatory actions during development.

The GRN was built around TFs and their ability to activate and repress a set of targeted genes, also known as their *regulons*. To identify how TFs structured the GRN, we evaluated the overlap between the regulons of each pair of TFs across the network and measured how similar or opposite was their inferred actions on shared targets (**Fig S4**). This delineated an *interactome* of TFs, identifying for each TF a set of preferential collaborators and competitors that tended to regulate genes through either identical or separates enhancers. Strongest interactions occurred not only between TFs with closely related binding motifs families (driven in part by similarity in targeted enhancer sites), but also across major brain regulators which are known to coregulate genes expression (**Fig. S4A**). For instance, the interactome of the TFs EOMES revealed a preferential collaboration with NEUROG2, NEUROD2 or MEF2C and a competition with FOXB1, LHX6 or ZNF823 across several targets in the GRN. Other examples included FOXP2 collaboration with ZNF548 and competition with TEAD4; or TBR1 collaboration with NFIB and ESR2 and competition with ZNF823 (**Fig. S4B**). Altogether, the built GRN integrated enhancer identification with gene expression to chart the regulatory effects and interactions of TFs in our organoid model (see GRN in **Table S3**).

### 2. Identification of master regulators behind neurogenesis progression and cell type specification

To explore how the enhancer-centered GRN captured regulatory events of human neural development, we next sought to identify the regulatory relationships central to the progression of neurogenesis. The GRN was therefore subset to only include genes differentially expressed between TD30/60 and TD0 (i.e., late vs early, termed timeDEGs) (**Fig. S5A-B, Table S4, T1**). Regulatory links in the GRN were then subset to include only links where change in expression of upstream TF was susceptible to drive differential expression of the downstream gene (**Fig. 2A, Methods**). This time transition subnetwork revealed two antagonizing groups of TFs driving major expression changes during early development (**Fig. 2A, Fig. S6C**). To further rank TFs’ abilities to drive expression changes, we computed for each TF an *activity perturbation score* that measured how its regulon was differentially affected between 2 conditions based on GRN prediction. The perturbation score metric is detailed in **Fig. S6**, with positive, negative, and near-zero perturbation scores indicating respectively, an increased, decreased or unchanged activity of the TF between two tested conditions. Applied to timeDEG, this perturbation analysis revealed that increased activity of TBR1, FOXP2 and POU6F1 and decreased activity of DMRTA1, FOSL1, and TEAD4 were driving gene expression changes across this time transition (**Fig. 2B**). Interestingly, some TFs with low change in expression level across time points had high activity perturbation scores (for instance ZNF490, SP1 or FOXN3). Such cases could stem from differences in posttranscriptional modulation of the TF, influencing for instance its DNA- or protein-binding abilities, although this could also result from expression changes in TFs acting on closely related regulons. The TD30 to TD60 transition, involving primarily an increase in neurogenesis, was driven by increased activity of TFs such as BHLHE22, NEUROD2, ESRRA or BHLHE40 (**Fig. S6D,E**).

**Figure 2.**
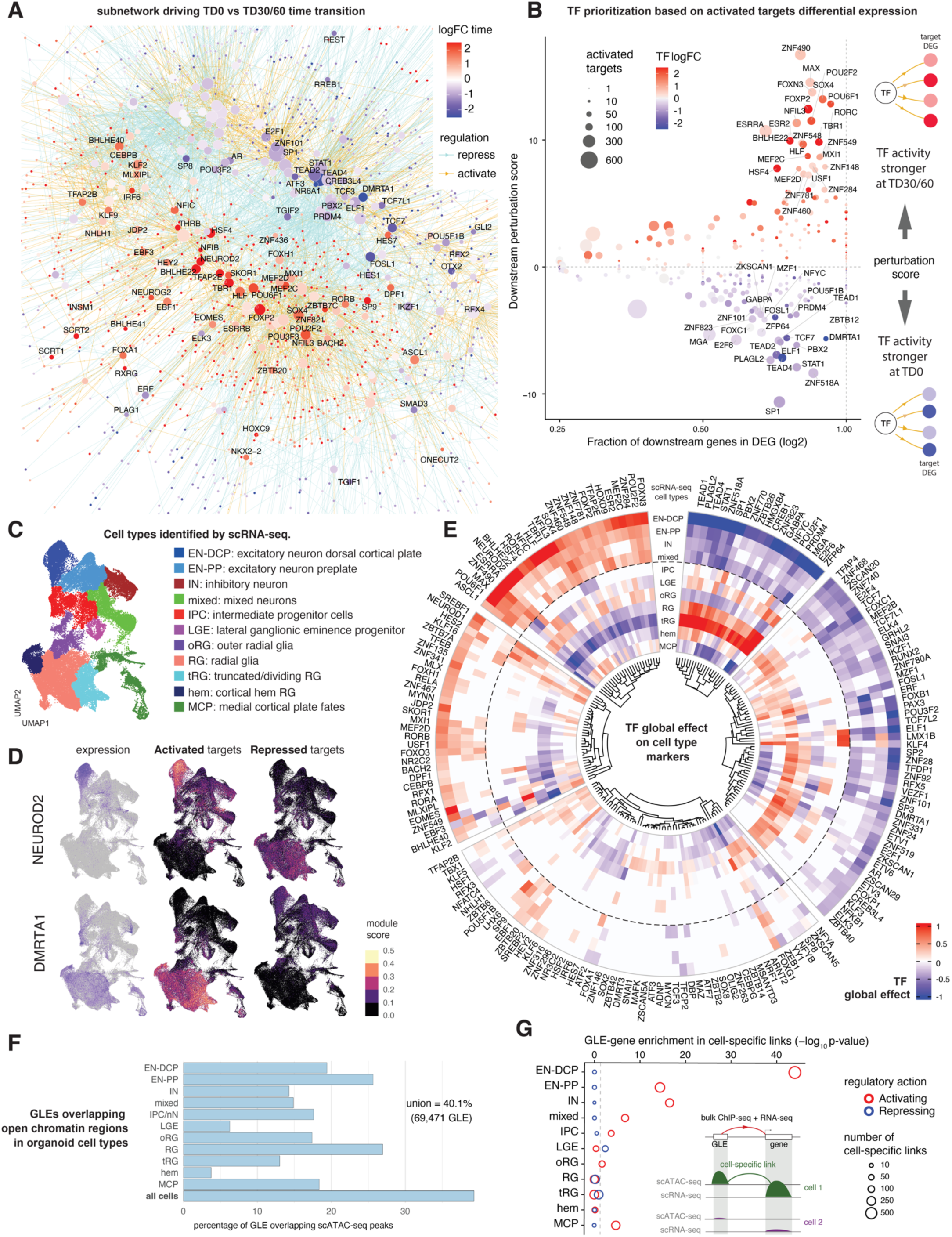
Master regulators driving neurogenesis and cellular diversity in forebrain organoids. **A.** Time transition subnetwork built from the GRN by retaining significant timeDEGs (TD30+60 vs TD0, |log_2_FC| > 0.25 and FDR < 0.01) and concordant links connecting them. Activating edges (yellow) need to connect timeDEGs with same direction of change and repressive edge (cyan) timeDEGs with opposite directions of change. Top TFs ranked by logFC are named. Kamada-Kawai layout with distances proportional to the direction and strength of the regulation as in **Fig. 1G**. **B.** Dot plot showing the fraction of TF’s targets that are timeDEG (x-axis) and the TF activity perturbation score (y-axis, schematic). Dot color shows the TF’s own log_2_FC in timeDEG. Dot size=regulon size in the GRN. The perturbation score ranks each TF’s ability to drive the expression changes of its regulon (here, across time) and is computed from GRN prediction and DEG results for the downstream target only. As shown by yellow arrows in right side schematic, only activating relationship were used for the perturbation score in this plot. Top 25 TF by activity perturbation score are named (see **Methods** and **Fig. S6**). **C.** UMAP plot colored by cell types identified in 72 scRNA-seq samples from TD0, 30 and 60 organoids ^15^ of the same individuals used for this study. **D.** UMAP plots colored by expression level for the TF NEUROD2 and DMRTA1 (left panels, blue=high, grey=low). To infer if TF’s targets also presented cell-type specific expression, the respective downstream “activated targets” and “repressed targets” of each TF’s regulon were used to derive a module expression score in the scRNA-seq data and positive module scores per cell are plotted in the UMAP (see also **Fig. S5C**). **E.** Circular heatmap showing the global effect of each TF on cell type-specific gene expression quantified by scRNA-seq (see **Data S5** for cell type markers). Results for each cell type plotted as concentric circles. TF’s global effect is a perturbation score applied to cell-type DEGs (as in **B, Methods**). Results are grouped by k-mean clustering (k=5) and hierarchical clustering (Euclidean distance, complete method). **F.** Fraction of all GLEs that intersect with scATAC-seq peaks in organoids (overlap with peaks identified in at least 3 out of 29 scATAC-seq samples, see **Fig. S5** for scATAC-seq peaks overlap and QC results). “all cells” = peaks called using all cells. Cell type annotations were identified by label transfer using scRNA-seq from **C** as reference. **G.** Enrichment of cell type-specific regulatory relationships (cell-spe.links) among all enhancer-gene regulatory relationships identified (**Table S2,T3**). *Cell-specific links* were defined as cases where the enhancer overlaps with scATAC-seq peaks (>= 3 samples) and the linked gene is a cell type-specific marker of the same cell type. Enrichment was calculated separately for activating and repressing enhancer-gene links (red and blue circles, respectively). p-value of one-tail Fisher exact test reported.

Organoids develop from aggregates of neural progenitors that generate a diversity of neurons over time^3,26^. Using scRNA-seq data, we have shown that our organoid method generated a diversity of forebrain neuron fates (**Fig. 2C**) and that the proportion of different cell types tended to vary between organoids from different individuals ^15^. Since bulk-level gene expression in organoids is strongly driven by changes in cell type composition, we hypothesized that the GRN captured in part the regulatory events driving variations in cell fates. Indeed, we observed that many timeDEGs had cell type-specific expression in scRNA-seq data (**Fig. 2D, Fig. S5A-C**), including many TFs drivers of time transition identified from the GRN (**Fig. 2B**). Coherently, TFs’ activated regulons tended to be co-expressed in the same cells as their upstream TF, while repressed part of their regulon had often inverted expression pattern across cells (**Fig. 2D, Fig. S5C**). This suggested that many central TFs in the GRN are driving binary choices between cell type identities or states. We therefore applied the perturbation activity score metric to cell-type specific DEGs (**Table S5**) to evaluate each TF’s capacity to drive cell type-specific expression pattern and identify putative activators and repressors of specific cell fates (**Fig. 2E**). As expected, many TFs controlled the distinction between neurons and progenitor cells (e.g., FOXP2, SOX4, TFAP2E) upregulating general neuron fate and repressing radial glia, while some TFs were preferentially regulating genes expressed by a particular neuronal subtype. Among the latter, EOMES, BHLHE22, NEUROD2, and NFIC were the strongest activators of excitatory neurons of the dorsal cortical plate (EN-DCP), while E2F6, TCF7L2 and POU2F1 were the strongest repressors of that fate, consistent with mouse loss-of-function studies ^27,28^.

To evaluate how cell type-specific regulatory events were captured in data derived from bulk-level ChIP-seq, we intersected our list of GLEs with open chromatin peaks called from 29 scATAC-seq samples collected from the same organoid preparations (**Table S2**). We found that 40.1% of GLEs overlapped with an ATAC-seq peak identified in a cell type or shared across cells (**Fig. 2F**). Furthermore, activating GLE-to-gene regulatory links from the bulk GRN were enriched in cell type-specific links (i.e., cases where the GLE overlapped a scATAC-seq peak and the linked gene showed increased expression in the same cell type in scRNA-seq data), suggesting that the bulk-derived regulatory network captured cell type-specific regulatory events and increasing confidence in our characterized enhancer elements. This enrichment was more significant for neuronal cell types and particularly for EN-DCP (**Fig. 2G**). Altogether, these analyses suggested that identification of enhancers from bulk epigenome along the developmental timeline captured the complex regulatory events underlying the development of a multilineage system.

### 3. ASD probands show dysregulation in TF regulators of excitatory and inhibitory neuron specification

We have previously shown by scRNA-seq that organoids derived from macrocephalic ASD individuals in the present cohort (macroASD) showed an increase in EN-DCP cell proportions and gene expression as compared to their unaffected fathers, whereas normocephalic ASD individuals (normoASD) showed a diametrically opposite phenotype ^15^. Here, by analyzing paired ASD versus control-father DEGs by bulk RNA-seq (pDEG in **Fig. S7**) and differentially active enhancers (pDAE in **Fig. S8A**) we confirmed those opposite phenotypes at the bulk transcriptomic and epigenomic levels (n=4 macroASD pairs and n=5 normoASD pairs, analyzed at TD0 and at TD30 + TD60 (merged together to increase power), see **Methods** & **Table S4, T1-2**). As in the scRNA-seq dataset, most DEGs were changed either in a cohort-specific manner or in opposite directions, with only 13 concordant pDEGs between the macroASD and normoASD cohorts (out of 641 pDEGs in either cohort at TD30/60, **Fig. S7C**). Bulk transcriptome data reproduced the upregulation and downregulation of gene markers of EN-DCP cells in macroASD and normoASD, respectively (**Fig. S7B-D**). This was further supported by an upregulation of enhancers upstream of several of those EN-DCP genes in macroASD pDAE (including of *EMX1*, *NEUROG2*, *NFIA*, and *LHX2*) (**Fig. S8A,B)**. In addition to this EN phenotype, a strong downregulation in genes related to the inhibitory neurons (IN) lineage (e.g., *DLX*-genes and *GSX2*) was observed at both TD0 and TD30/60 in macroASD pDEG (**Fig. S7A-D**).

To deconvolute the regulatory drivers of ASD-associated changes, we identified the upstream TFs of macroASD pDEGs in the GRN (**Fig. 3A**). This **macroASD subnetwork** comprised 96 pDEGs (out of 420) controlled by 102 TFs, of which 19 were pDEGs themselves (**Fig. 3A**). We then applied TF activity perturbation analysis (**Fig. S6**) to systematically rank each TF’s ability to drive differential expression of its downstream targets in macroASD. This analysis highlighted an “EN regulatory hub” composed of the master regulators BHLHE22, EOMES and FOXG1 as the strongest drivers of the macroASD pDEG (**Fig. 3B**) which were all strongly associated with EN-DCP fates regulation (**Fig. 2E**). BHLHE22, EOMES and FOXG1 acted on several downstream TFs controlling EN-DCP fate such as EMX1, LHX2, ZBTB18 and BCL11A (**Fig. 2E, 3A,B**). While we had lower sensitivity to detect pDAEs than pDEGs, in several cases the intermediate enhancer connecting the TF to the downstream pDEG in the GRN was itself a pDAE (**Fig. S8B**). For instance, in macroASD, the upregulation of the TF *ZBTB18* could be associated with the upregulation of two of its enhancers (ID: 123747, 124166) themselves also bound by the TFs BHLHE22, TBR1 and TFAP2C, three upregulated pDEGs. Similar intermediate pDAEs linked FOXG1 to *BCL11A*, FOXG1 to *LHX2*, and EOMES to *EMX1* (**Fig. S8B**). Such cases identify potential “propagation paths” of the phenotype where TFs’ expression changes caused epigenomic changes in targeted enhancer elements and ultimately differential expression of secondary downstream targets. In agreement with the observed decreased in GABAergic interneuron (IN) genes in macroDEGs, the known inhibitory lineage TFs ASCL1 and LHX6 ranked among the top driver TF with decreased activity in macroASD (**Fig. 3B**).

**Figure 3.**
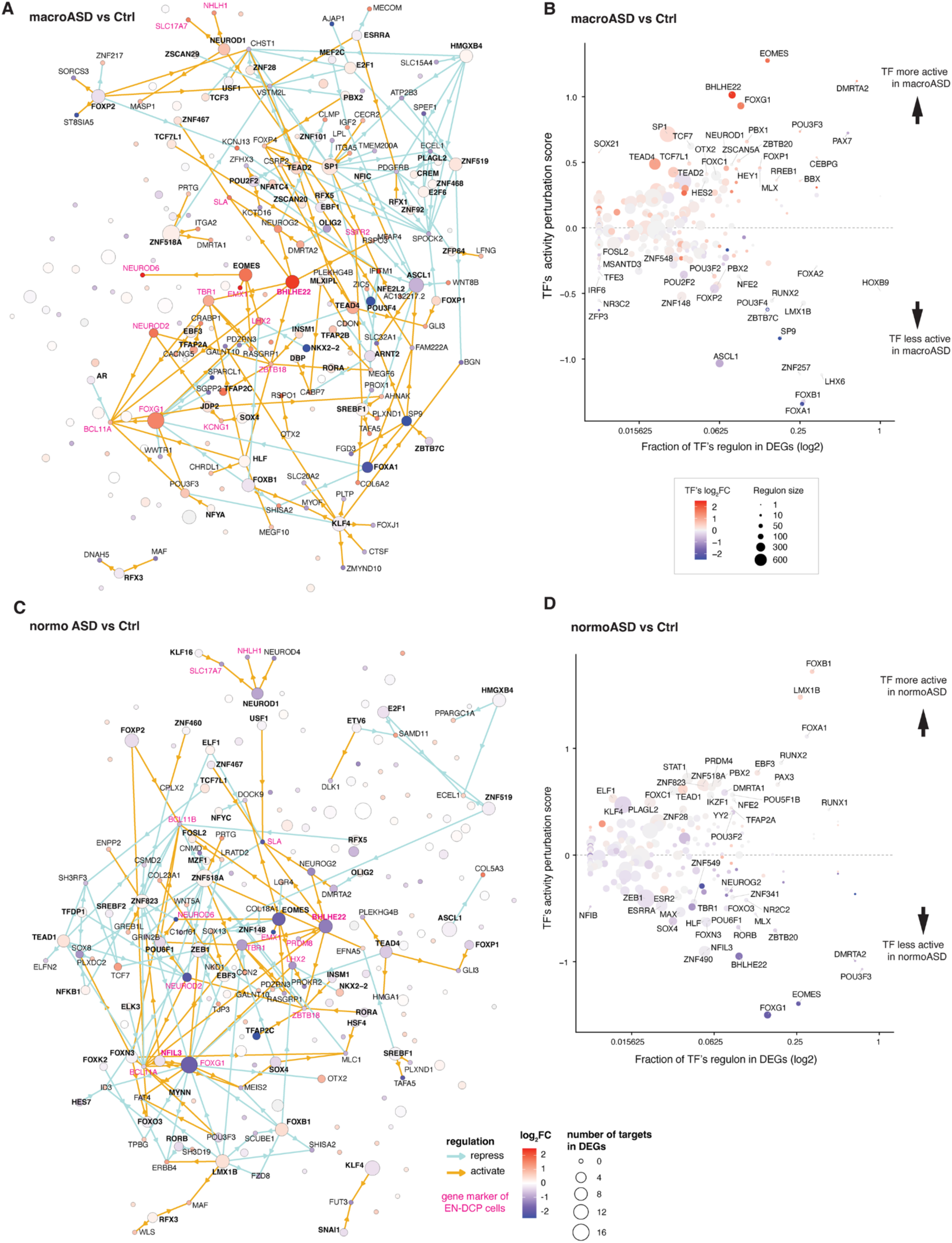
ASD probands have head-size cohort-specific alterations of the regulatory network coordinated by central transcription factors like FOXG1. **A,C.** TF-to-gene GRN subnetwork depicting pDEGs (paired DEGs between ASDs and control fathers using TD30 and TD60 bulk RNA-seq expression level) and their upstream TFs for macroASD (**A**) and normoASD (**C**). Color of the genes represents expression change, while color of arrows represents the regulatory action (activation/repression) from the GRN (**Fig. 1G**). Genes are disposed identically in **A** and **C**. Dot size show number of targets in the resulting network. Gene markers of EN-DCP cells from scRNA-seq data are in red. **B,D.** TF ranking based on their activity perturbation scores against macroASD pDEGs (**B**) and normoASD pDEGs (**D**). TFs are plotted based on the fraction of their downstream targets that are significantly pDEGs (x-axis) versus the TF’s activity perturbation scores (y-axis) which evaluates each TF based on how strongly its predicted effect in the GRN (i.e., activation/repression) accounts for the observed pDEGs in its regulon. Dot colored is the log2FC value of the TF itself (ASD vs Ctrl), dot size= regulon size.

Similarly, the **normoASD subnetwork** placed 85 pDEGs (out of 315 pDEG at TD30/60) under the control of 103 upstream causative TFs. In normoASD, the “EN hub” composed of the master regulators BHLHE22, EOMES and FOXG1 showed strong decrease in activity, causing a secondary downregulation of *EMX1*, *LXH2, NEUROG2, TBR1, ZBTB18, BCL11A,* and *NEUROD6* (**Fig. 3C,D**). These TFs often co-regulated each other, with, for instance, LHX2 downstream of EOMES and FOXG1, and ZBTB18 controlled by both BHLHE22 and TBR1. Interestingly, the top three TFs with increased activity in normoASD (i.e., LMX1B, FOXB1 and FOXA1, **Fig. 3D**) were direct repressors of FOXG1 and were affected in an opposite way in the macroASD cohort. Taken together, the GRN identifies opposite dysregulation of two triads of co-regulated TFs (BHLHE22, EOMES and FOXG1 against LMX1B, FOXB1 and FOXA1) as the strongest drivers of the opposite transcriptomic phenotypes observed in our macro- and normo-ASD cohorts. Those results reinforce that organoids from these cohorts of ASD probands revealed divergent transcriptomes as well as epigenomes.

To expand our study beyond ASD idiopathic individuals, we performed a similar analysis to explore the regulation and interconnection of genes associated to rare variants in ASD (SFARI genes). The built GRN can allow to understand how genes carrying rare ASD mutations interact together to perturb brain development. For instance, using the list of identified enhancers and their upstream networks (**Data S2**), we could identify the upstream enhancers and TFs that regulate the SFARI risk genes CHD3 (**Fig. 4A**). Associating upstream regions de facto extends the genomic regions of interest for the identification of disruptive variants of any given risk gene to noncoding regions or indirectly associated coding regions. The GRN could also be used to identify paths of downstream regulatory convergence between ASD risk genes. We identified regulatory relationships associating 348 ASD-associated risk genes identified in SFARI (out of ∼1000) (**Fig. 4B**). Risk genes were preferentially associated in the GRN (Fisher exact test two-sided p-value = 0.0015), notably because they included 33 highly interconnected TFs. Some TF risk genes such as BCL11A were downstream of important TF such as FOXP2, TBR1 or FOXG1, which are part of the EN regulatory hub implicated in the idiopathic ASD subnetwork (**Fig. 3A**).

**Figure 4.**
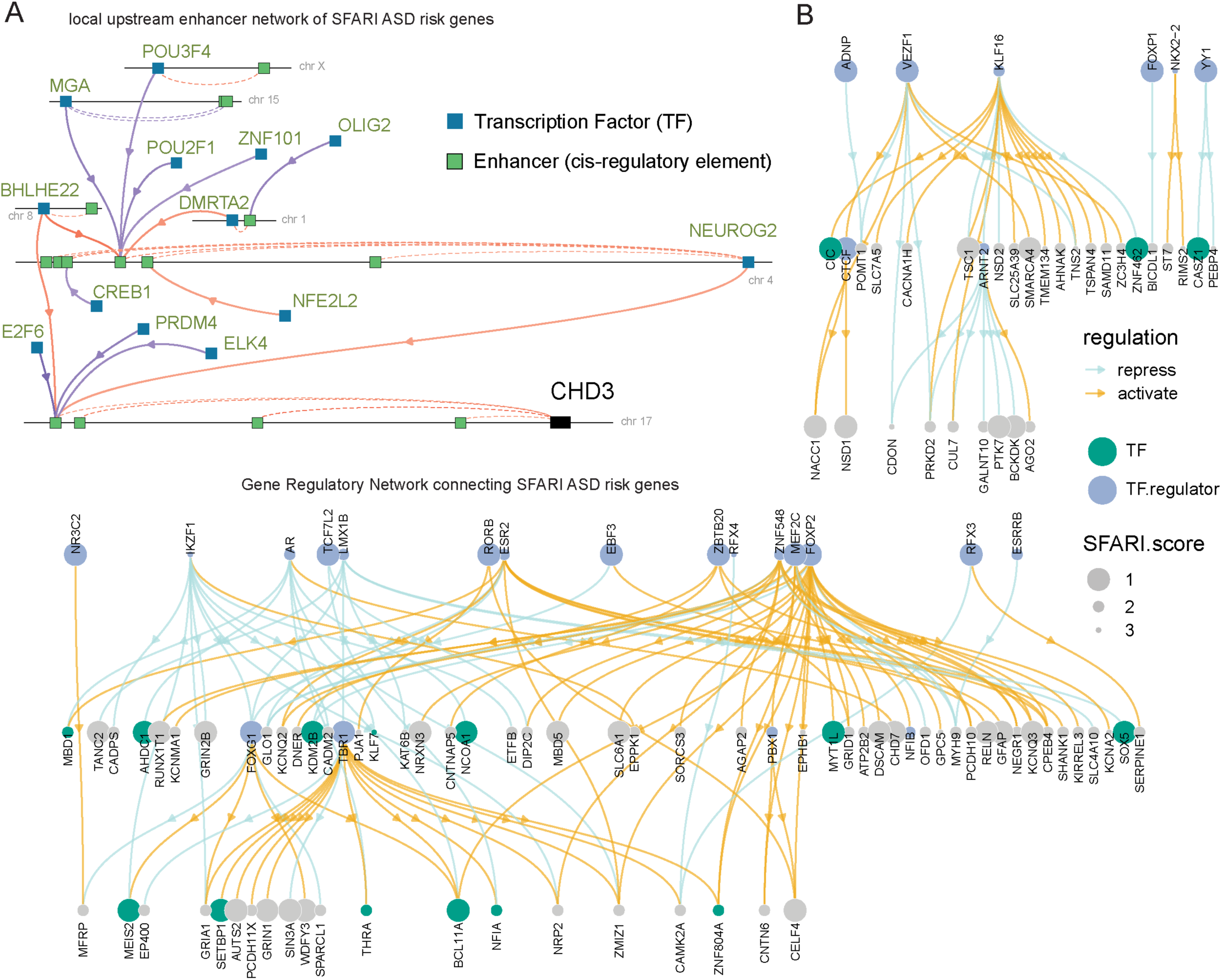
regulation and interconnection of known ASD risk genes. **A.** Example of upstream enhancer subnetwork for the ASD risk gene and chromatin modifier CHD3. Enhancers (green boxes) are connected to their downstream genes by dashed lines. TF (blue boxes) are connected to their downstream enhancer targets by arrows (red=activation, blue=repression). When in turn TF have themselves an enhancer controlling their expression, it is indicated on the linear genome. Enhancers are positioned at scale in the linear genome (their width is not represented, chromosome number is indicated, see **Data S2**,T4 for positions). Note the 6 degrees-separation between CHD3 expression and the TF OLIG2. **B.** GRN restricted to links connecting 348 ASD risk genes (risk score indicated by size, with score=1 more confidently associated to ASD as defined by SFARI database).

### 4. Enhancer cross-validation using CRISPRi

To validate our ability to identify enhancers using ChIP-seq and functionally confirm their downstream targets, we applied CRISPR interference (CRISPRi) to our organoid system (**Fig. 5A**). We used a nuclease-deactivated dCas9 fused to the Zim3 KRAB (Krüppel-associated box) domain, which recruits endogenous factors that trimethylates histone H3 on Lys9 (H3K9me3), silencing the activity of targeted regulatory element ^29,30^. We generated an iPSC line with constitutive expression of Zim3-*dCas9*, a mScarlet reporter, and a hygromycin selection cassette by PiggyBac transfection of the iF53 vector in a male control iPSC line (RDH913-03#9) followed by hygromycin selection (**Figure S9A; Methods)**.

**Figure 5.**
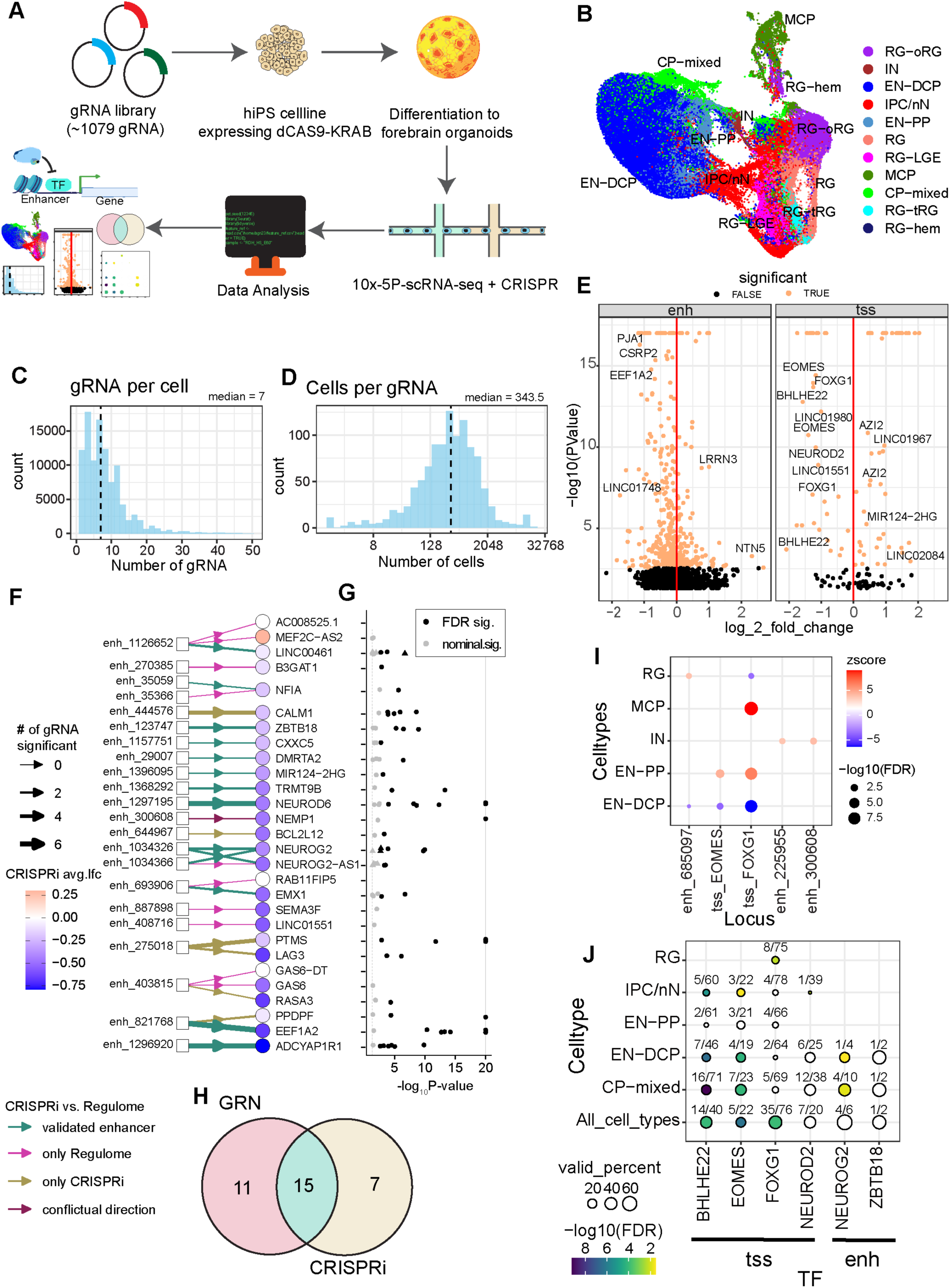
CRISPRi validation of GRN enhancers. See also Figure S9. **A.** Workflow representing enhancer validation by a pooled CRISPRi approach. **B.** UMAP at TD30 annotated by cell types as in Fig.2C. **C-D.** Histogram showing distribution of the number of gRNAs per cell (C) and cells per gRNA (D) at TD30. **E.** Volcano plots displaying gene expression change in gRNA-expressing versus non-expressing cells for genes located in ±1MB region centered on the targeted locus; gRNAs are separated by promoter-gRNA (tss) and enhancer gRNA (enh). Orange = significant differential expression with FDR < 0.1. **F.** Regulatory graph comparing enhancer-gene links (boxes-to-circles arrows) identified by CRISPRi to links identified in the GRN (arrow color, with “validated” (green) when significant in both experiments). Circle color=CRISPRi-averaged log2-fold change, arrow thickness show number of significant gRNA. **G.** Dot plot shows p-value for each gRNA perturbing the corresponding gene expression (black = FDR <0.1). **H.** Venn Diagram showing intersection of GLE-gene links predicted by GRN and links called by CRISPRi experiment. Intersection corresponds to 46.8% of total links tested. Link with conflicting result was added in both sets for representation. **I.** Significant change in cell fate due to perturbation of promoter (tss) or enhancer (enh_ID#, see **table S1** for precise region). color= z-score compared to overall distribution; dot size = significance (FDR-corrected z-test) (**Methods**). **J.** Validation rate of predicted downstream targets of TF after gRNA mediated silencing of TF’s expression (dot size = percent of GRN-targets validated with number of genes indicated (validated / tested), color= Fisher exact-test FDR corrected significance for enrichment of GRN-targets in CRISPRi results).

We constructed a gRNA lentiviral library to target 104 loci of interest, selecting multiple gRNAs per target locus ^31,32^, **Methods**). This included 94 putative enhancer loci (enh-gRNA; median 10 gRNAs per locus, range 6-10) selected from our study among the 173,437 GLEs initially identified (**Fig. 1B, Table S2**). It also included gRNAs targeting the promoter regions (prom-gRNA) of 6 TFs (*BHLHE22, FOXG1, EOMES, NEUROD2, LHX6, LXM1B;* 10 gRNAs per gene) and 4 positive control genes (*GRN, CDH2, TFRC, UBQLN2;* 3 gRNAs per gene) previously shown to be efficiently repressed by CRISPRi in iPSC-derived neurons ^33^ to evaluate the efficacy of CRISPRi. The library included ∼10% of non-targeting gRNAs as negative controls (scrambled sequences), for a total of 1079 gRNAs **(Table S6**). This gRNA library was transduced by lentivirus into the Zim3-dCas9 iPSC line at high multiplicity of integration (MOI of ∼5), as CRISPRi can be efficiently multiplexed to test multiple gRNAs per cell to reduce cost^34^. After puromycin selection, the gRNAs+Zim3-dCas9 line was differentiated into forebrain organoids and cells were analyzed by scRNA-seq at TD0 (∼10k cells) to validate the quality of the preparation. The majority of cells mapped to cell types previously described at this stage ^15^, suggesting that the Zim3-dCas9 and gRNA lentiviral library integration did not alter the capability of the RDH913-03#9 iPSC line to differentiate toward forebrain (**Figure S9B,C,D**). The downregulation of positive control genes and *FOXG1* in the respective gRNA-carrying cells demonstrated that the Zim3-dCas9 system efficiently silenced targeted chromatin regions and decreased gene expression (**Fig. S9E**).

The perturbation of targeted enhancers and TFs was then assessed at TD30 with high cell coverage (6 scRNA-seq libraries, with a total of ∼103k cells passing QC). After unsupervised clustering, cells were mapped to our scRNA-seq reference ^15^ identifying forebrain RG and its subtypes, IPC/nN, EN-DCP, IN, and medial cortical plate (MCP) cells (**Fig. 5B and S9F**). Cells contained a median of 7 gRNAs per cell (**Fig. 5C**) and ∼95% of the library’s gRNAs were represented in the cell population at various frequencies, with a median of 343.5 cells per gRNA (**Fig. 5D**). For each gRNA, differential expression was tested for genes in a 1Mb window around the targeted locus using SCEPTRE ^35^ (see **Methods**) comparing cells expressing the gRNA to all other cells (**Fig. 5E, S9G,H**). Genes downregulated by at least 2 independent gRNAs targeting the same enhancer were considered as validated cis-activated genes targeted by that enhancer (with differential expression meeting FDR<0.1 significance, and no gRNA with significant discordant upregulation). We excluded cases where the gRNA was located inside the downregulated gene body as gRNA-Cas9 binding could have directly disrupted Pol2 elongation. This conservative analysis identified at least one downregulated target gene for 69 out of 94 of the putative enhancer loci tested (**Table S6**), validating the presence of an enhancer element at those GLE loci initially identified in our analysis.

To compare the results of CRISPRi to GLE-gene regulatory links identified during GRN construction based on correlation of enhancer activity with expression of downstream gene(s) (**Fig. 1**), we focused on 22 GLE loci with significant downstream gene(s) identified in both experiments, comparing 32 GLE-gene links tested in both experiments (**Fig 5F-G, Methods**). Among those, 15 GLE-gene activating links in the GRN showed concordant results in CRISPRi (validated regulatory link), corresponding to a 46.8% intersection rate between tested enhancers across the two experiments (**Fig. 5H**). One link was found to have opposite direction of regulation between the two experiments, while the remaining 17 links were called only by one approach. Validated loci included enhancers activating important genes for EN-DCP neurogenesis including *NEUROD6*, *EMX1*, *NEUROG2* and *ZBTB18*. Aside from identifying downstream targets of targeted enhancers, we also identified 5 loci for which the gRNA presence induced shifts in cell composition, including the promoters of *FOXG1* and *EOMES* (**Fig. 5I**), supporting that the perturbed genes were involved in organoid patterning, lineage definition, and/or neurogenesis. The disruption of *FOXG1* expression created a major shift towards medial and preplate cell types (**Fig S9I**). *EOMES* disruption appeared to redirect cells from EN-DCP to EN-PP, concordant with the role of EOMES in indirect neurogenesis and IPC function in mouse studies^36^. To identify potential downstream targets of TF perturbed by CRISPRi, we merged all the gRNAs affecting the same TF (promoter-gRNA or enhancer-gRNA affecting gene expression) and tested for genome-wide differential expression between cells with or without the tested gRNA, globally or at the cell type level (**Methods**). Although this analysis cannot parse between direct and secondary effect of the CRISPRi-induced TF’s downregulation and focused on genes expressed in this particular experiment only, it validated several downstream targets of EOMES, BHLHE22, as well as targets identified for ZBTB18 and NEUROG2 in the GRN (**Fig. 5J**). For FOXG1, out of 111 GRN-predicted FOXG1 targets, 76 were expressed in the CRISPRi assay, of which 35 were significantly differentially expressed, resulting in a validation rate of 46%.

To our knowledge, this is the first CRISPRi screen in forebrain organoids. While cost and technical limitations make screening larger gRNA libraries challenging, the successful CRISPRi perturbation of enhancers and promoter loci allowed us to functionally validate several components of the GRN, reinforcing their involvement in cortical excitatory neurogenesis.

### 5. FOXG1 loss of function validated the gene regulatory network

FOXG1 was identified as one of the top candidate drivers of the ASD phenotypes in the GRN, so we did an exhaustive analysis of its central influence on the development of forebrain organoids. *FOXG1* upstream regulatory network comprised 14 activating GLEs in its *cis*-regulatory region (chr14:28,751,330–30,3825,86), which are bound by 11 repressors (including the top 3 TFs with increased activity in normoASD, FOXB1, LMX1B, and FOXA1) and 2 activators (NFIL3, ZEB1) with additional other indirect putative regulators upstream (**Fig. 6A**). In scRNA-seq data, *FOXG1* was strongly expressed in forebrain RG cells, silenced in hem-RG, preplate (EN-PP) and medial cortical plate (MCP), and re-expressed in postmitotic EN-DCP and IN of the dorsal cortical plate (DCP) (**Fig. 6B**). Consistent with this, we found that several TFs that repressed *FOXG1* (such as EBF3, FOXB1, FOXA1, ZNF823 and LMX1B, **Fig. 6A**) activated EN-PP genes and/or repressed EN-DCP genes (**Fig. 2E**). Downstream binding targets of FOXG1 included 418 activated GLEs and 281 repressed GLEs across the genome, suggesting that FOXG1 could act as an activator or as a repressor of enhancer activity (**Fig. S10A, Table S2**). Through those enhancers, we identified 111 downstream genes in the regulon of FOXG1, including 8 TFs (*POU3F3, BCL11A, LHX2, MEIS2, MEOX1, OTX2, SOX13, ZNF503*, **Fig. S10B, Fig. S3**). FOXG1 activated targets were expressed in RG progenitor, IN (e.g., *MEIS2*) and EN-DCP (e.g., *BCL11A, LHX2*), while some of its repressed targets (e.g., *OTX2*) were expressed in cells with medial fates (hem-RG, and MCP cells) (**Fig. 6B, Fig. S10C**). Finally, inside the GRN, FOXG1 competes preferentially with FOXB1, LMX1B or POU5F1B and collaborates more specifically with FOXK2, POU3F3 or FOXO3 across shared targets (**Fig. S10D**).

**Figure 6:**
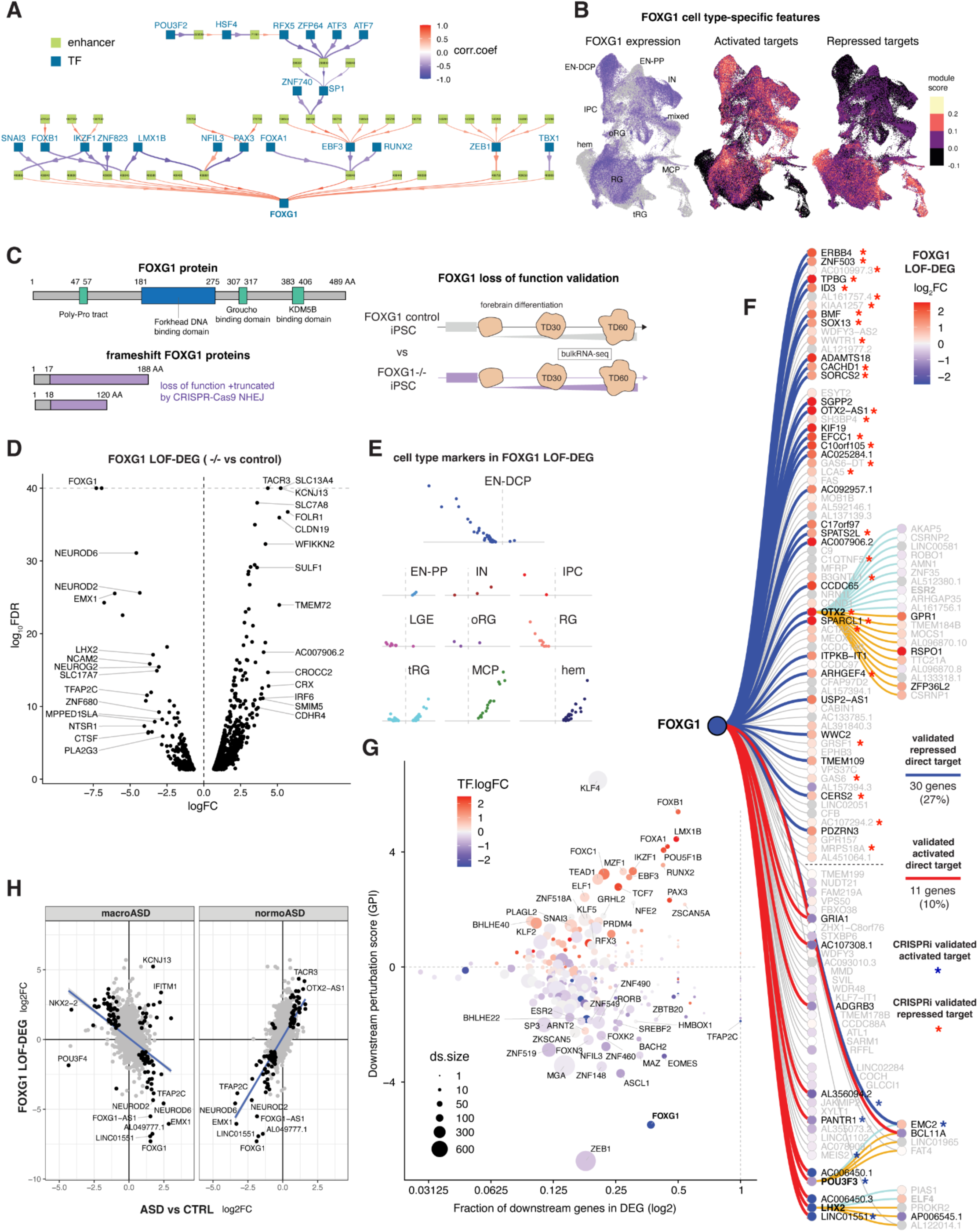
Validating the downstream targets of the master regulator FOXG1 by CRISPR loss of function. **A.** Upstream regulators of FOXG1 including their respective cis-GLE and upstream TFs. **B.** UMAP plots showing FOXG1 expression level in cell types juxtaposed with the positive module score expression of its downstream activated and repressed target genes. **C.** CRISPR-induced FOXG1 loss of function experiment. Schematic representation of the normal FOXG1 protein and the two predicted out-of-frame truncated proteins generated by CRISPR/Cas9 cleavage and non-homologous end joining (NHEJ), retaining only 17-18 amino acids (AA) of the normal FOXG1 protein and depleted of all the FOXG1-functional domains, including the forkhead DNA-binding domain. **D.** Volcano plot of FOXG1 LOF-DEGs (comparing FOXG1 LOF vs non-edited and the founder iPSC line). **E.** Volcano plots as in **d** highlighting FOXG1 LOF-DEGs that have cell type-specific expression, highlighting the strong downregulation of EN-DCP genes and upregulation of medial fate (Hem, MCP). **F.** Graph of FOXG1 regulon with targets colored by FOXG1 LOF-DEG results. Edges connecting FOXG1 to its targets are colored in blue for repression or red for activation when validated by FOXG1 LOF-DEG. Fisher’s exact test confirmed that both FOXG1 activated targets are enriched in genes downregulated during FOXG1-loss-of-function (p-value = 1.5 x 10^-6^) and FOXG1 repressed targets are enriched in genes upregulated during FOXG1 loss-of-function (p-value= 2.1 x 10^-13^). Direct target genes are organized from top to bottom or bottom to top based on their original correlation level with FOXG1 in the GRN. Asterisks indicate targets validated by CRISPRi at FOXG1 promoter (see **Fig. 5J**); note they are among the strongest correlated genes. Secondary targets of FOXG1 are also reported (through OTX2, POU3F3 or LHX2 regulation). **G.** Perturbation plot for FOXG1 LOF-DEG, plotting for each TF in the GRN, the fraction of its regulon in LOF-DEGs (x-axis) versus its activity perturbation score (as in **Fig. S6A**). Log-2 fold change of the TF itself (color) and its regulon size (dot size) are indicated. Note that this analysis uses the GRN and the FOXG1 LOF-DEG result to *naively* re-identify FOXG1 as one of the main drivers of the difference. **H.** Confusion plots comparing logFC results between FOXG1 LOF-DEG and ASD vs Ctrl pDEG for macroASD and normoASD. Top genes by distance from the origin are shown. Linear regression line is shown in blue (Pearson’s correlation R=-0.4 and 0.63, respectively, with both p-val < 2.2.10^-16^)

Several targets of FOXG1 were validated by CRISPRi (**Fig. 5J**). To further validate the regulon of FOXG1 in a focused experiment, we generated iPSC lines with FOXG1 loss of function mutations (FOXG1-/). CRISPR/Cas9 genome editing was used to introduce frame-shifting indels (see **Methods**), resulting in generation of a premature stop codon and a truncated FOXG1 protein lacking the forkhead-DNA binding domain (**Fig. 6C**). We then assessed differences in gene expression in organoids by bulk RNA-seq at TD30/60 by comparing two homozygous FOXG1-/-edited lines to two FOXG1+/+ control lines (termed LOF-DEGs, **Table S4,T3, Methods**). LOF-DEGs comprised 250 downregulated genes and 620 upregulated genes (**Fig. 6D**). The top downregulated LOF-DEG was *FOXG1* itself, suggesting that the FOXG1 exerts a positive autoregulatory effect on its expression or that the *FOXG1* truncated mRNA undergoes non-sense mediated decay. We observed that genes expressed in EN-DCP and RG were downregulated in LOF-DEG, while gene expressed in EN-PP, RG-hem and MCP, such as OTX2, were upregulated (**Fig. 6D,E**). This was concordant with the results of the CRISPRi-mediated *FOXG1* downregulation which showed that cells with gRNA targeting the *FOXG1* promoter exhibited a fate shift towards MCP and EN-PP (**Fig. 5I and S9I**). We also noted that cell cycle genes were overall downregulated (GO:0007049, p-value = 4.5×10^-11^ in FOG1 LOF downDEG, including genes such as MKI67 or TOP2A; **Table S4,T3**), which is consistent with microcephaly phenotypes resulting from FOXG1 LOF in humans^37,38^.

When comparing the FOXG1 LOF-DEGs to the GRN, 37% of the 111 FOXG1 targets predicted by the GRN (30 repressed and 11 activated) were significant LOF-DEGs with concordant direction of change (**Fig. 6F**). FOXG1 activated targets in the GRN were enriched in genes downregulated in FOXG1-loss-of-function (Fisher’s exact test p-value = 1.5 x 10^-6^) whereas FOXG1 repressed targets in the GRN were enriched in genes upregulated in FOXG1 LOF (p-value= 2.1 x 10^-13^) (**Fig. S10E**). For the remaining 70 FOXG1 targets in the GRN the direction of change in FOXG1 LOF-DEG remained concordant with the GRN prediction in a majority of cases (FDR > 0.05). Out of 41 gene targets validated by the LOF approach, 20 targets were also validated by CRISPRi, including *ERBB4, OTX2, POU3F3, MEIS2, SOX13* and *ID3* (**Fig. 6F**). Next, the LOF-DEG results were used as an input for TF activity perturbation analysis, which yielded FOXG1 as one of the strongest drivers of the LOF-DEGs with predicted decrease in activity (**Fig. 6G**), confirming our approach for identifying driver TFs from differential expressions using the GRN (**Fig S6**). Thus overall, the FOXG1 LOF experiment validated the constructed GRN. Finally, we observed that gene expression alterations induced by the FOXG1 LOF experiment were negatively correlated with macroASD pDEGs and positively correlated with normoASD pDEGs (**Fig. 6H**). This was attributable to top upregulated genes in macroASD being downregulated in LOF-DEGs, and to upregulated genes in normoASD being upregulated in LOF-DEGs. This similarity between ASD pDEGs and FOXG1 LOF-DEGs supported the hypothesis that a FOXG1 dysregulation in opposite directions drives a strong part of the ASD phenotype in each cohort.

Altogether, this is consistent with the role of FOXG1 in repressing medial fates in mammalian forebrain development ^39,40^, where its inactivation leads to a decrease of dorsal cortical fates and an increase in medial fates. Put together, the forebrain organoid system confirmed and reproduced the fundamental role of FOXG1 during development.

## Discussion

Using parallel epigenomic and transcriptomic datasets we constructed a regulome encompassing gene and enhancers in developing organoids. We leveraged the identification of proximal and distal gene-enhancer relationships and TFBS to infer regulatory action of TF on both enhancer activity and linked gene expression. The obtained network captured the driving forces behind time transitions and cell type specification during neurogenesis. Experimental validation via CRISPRi and FOXG1-LOF analyses strengthened the biological relevance of the network and highlighted specific TF cascades, notably implicating FOXG1, BHLHE22, and EOMES in the regulation of excitatory neuron specification in ASD.

By using iPSCs derived from idiopathic ASD individuals we identified the TFs and dysregulated enhancers that drive the imbalance of excitatory neurons previously discovered in this ASD cohort ^15^. Specifically, we suggest that a dysregulation of FOXG1, BHLHE22 and EOMES expression in opposite directions in ASD with macrocephaly and ASD with normocephaly drives the excitatory neuron imbalance in opposite directions in these ASD subtypes. Dysregulated genes in the FOXG1 LOF experiment were largely overlapping with those affected in macroASD and normoASD, strongly suggesting that FOXG1 is a central causal factor of differential gene expression in ASD. While based on a limited number of individuals, our study is a first draft of gene regulatory interactions disrupted in this disorder during early neural development.

A limitation of existing GRNs is the reliance on correlation analyses to define regulatory links. In our case we limited the use of correlation and used it to supplement primary signals such as noisy predictions of TFBS, although we acknowledge that TF transcript levels are not always predictive of TF activity. Yet, GRN demonstrates considerable predictive power as evidenced by the CRISPRi and FOXG1 LOF validations, the moderate validation rate for enhancer-gene links suggests that additional layers of regulatory control such as chromatin conformation changes, combinatorial TF binding, and context-dependent enhancer activity are not fully captured by our current model. Interestingly, while the great majority of enhancers were activators of gene expression, negative correlations between TF levels and enhancer activity were just as abundant as positive one, suggesting that ∼50% of developmental TF could act as repressors and confirming the notion that repression is essential for establishing precise patterns of gene expression during development ^41^. Future GRN could incorporate proteomic information to account for post-transcriptional and post-translational modifications as important determinants of TF activity. While we could delineate from our GRN major collaborators and competitors of each TF (**Fig. S4**), further data collection would also help refine non-linear combinatorial effects occurring when multiple TFs co-regulate the same enhancers and genes. Another limitation of our GRN approach was the reliance on bulk-level data. This precluded a clear understanding of whether observed links in the GRN were specific to a given cell type or shared across cells. Nonetheless, we intersected the GRN with our own as well as external scRNAseq and scATAC data (**Figs. 1I and 2C-G**) and have found supporting evidence from single-cell data. Specifically, when genes are expressed in a cell type, their linked enhancers were also active in the same cell type, and such phenomenon involved active enhancers more often than repressing enhancers (**Fig. 2C-G**). While we show that the network captured major cell type specification, other subpopulation-specific regulatory interactions can be further delineated by expanding sample collections, including later stages and larger cohorts of individuals, and by incorporating single-cell level information in building of the network. Indeed, scATAC-seq data that captures open chromatin regions in each cell type overlapped only in part with ChIP-seq data and our regulome showed limited overlap with a recently described scATAC-seq organoid regulome^14^, suggesting that combining both histone marks and open chromatin data at single cell level could fully resolve the regulatory events occurring in each cell type.

Altogether, the regulome establishes a hierarchy between TFs and cognate enhancers that controls the dynamics of cell differentiation during typical and atypical human neurodevelopment. We show that fundamental TFs like FOXG1 can drive cascades that are extremely complex in humans, with hundreds of downstream enhancers and genes. Untangling the complexity of these regulatory interactions through regulatory network analysis may reveal the paths of convergence behind the genetic heterogeneity of human developmental disorders such as ASD, schizophrenia or intellectual disabilities.

## Supporting information

Suppplemental Figures

## Resource availability

All unpublished codes are described in the Methods section of the paper. This study did not generate new unique reagents or DNA constructs.

Primary cell lines and iPSC lines are shared via the Infinity BiologiX LLC repository (https://ibx.bio/) or available from the corresponding author after MTA.

Datasets reported in this study are available through the NIMH Data Archive (NDA). As specified in our consent form, data are available under controlled access for patient’s privacy.

The scRNA-seq and scATAC-seq data are under collections #C3957, url: https://nda.nih.gov/edit_collection.html?id=3957 and collection #2821, url: https://nda.nih.gov/edit_collection.html?id=2821

The bulk RNA-seq and bulk ChIP-seq data are under collection #C2424, url: https://nda.nih.gov/edit_collection.html?id=2424.

Data will be accessible through study #2322, url: https://nda.nih.gov/study.html?id=2322

## Acknowledgments

We are grateful to the families and children for their participation in this study. We thank Soraya Scuderi for help with the ChIP-seq protocol. We acknowledge Elise M. Cummings, Gloria Han, and Kelly Powell for help with subject recruitment and clinical characterization to produce iPSC lines and Arijit Panda for help with data upload into NDA and sharing. We thank Guilin Wang and Christopher Castaldi, and the Yale Center for Genome Analysis for library preparation, deep sequencing and Cell Ranger analysis. We thank Caihong Qiu and Jason Thomson at the Yale Stem Cell Center for the generation of the iPSC lines. We acknowledge the Yale Center for Clinical Investigation for clinical support in obtaining the biopsy specimens.

## Author contributions

Conceptualization, design, supervision: FMV, AAbyzov

Patient recruitment and acquisition of clinical data: McP, KP, PV, KC, LT

Skin biopsies: ASzekely

Primary cell culture, reprogramming and iPSC QC: LT

Organoid generation and processing for multiomic assays: JM, AJ, AS, Aamiri

Bulk RNA-seq and ChIP seq bioinformatic analysis: FW, SN, AJ

scATAC-seq bioinformatic analyses: FW

Zim3-dCas9 construct generation: MPN, DMR, MEW

CRISPRi experiment: AN, AJ, BL, CAG, GEC

gRNA library cloning, lentivirus production: MPN

FOXG1-loss of function lines experiment: AJ

Secondary bioinformatic analyses: AJ, SN, AN, DC

Writing—original draft: AJ

Writing—review & editing: AJ, AN, JM, FW, DC, AAbyzov, FMV

## STAR METHODS

### Lead contact

Further information and requests for resources and reagents should be directed to and will be fulfilled by the lead contact, Flora M. Vaccarino.

### Human Subjects/Ethical statement

All iPSC lines used in this study were described in a previous report where subject recruitment, clinical information and iPSC generation are fully described ^15^. Human subjects were recruited through several research projects at the Yale Child Study Center. Written informed consent was obtained from each participant enrolled in the study, and all research was approved by the Yale University Institutional Review Board (HIC# 1104008337) and Yale Center for Clinical Investigation at Yale University and was performed in accordance with the Declaration of Helsinki. Human participants’ names and other HIPAA identifiers were removed from all sections of the manuscript, including supplementary information. The participants agreed to data sharing of genomic unidentified data using controlled data access. All methods were performed according to relevant regulations and guidelines of Yale University, including the Biological Safety Committee and ESCRO committee of Yale University.

### Organoid preparation

All iPSC lines and forebrain organoids preparations used in this study were identical to a previous report (see **Data S1**) ^15^. Briefly, IPSC lines from controls and ASD-probands were propagated in parallel and grown in mTeSR1 media (StemCell Technologies). Forebrain organoids were generated using dual SMAD inhibition (SB431542, 10μM and LDN193189, 1μM, 7 days), followed by a proliferation phase (FGF2, 10 ng/ml + EGF, 10 ng/ml, 7days) and a terminal differentiation phase (starting at day 17, denominated TD0) in terminal differentiation media composed of Neurobasal medium supplemented with 1% N2, 2% B27 w/o vitamin A, 15 mM HEPES, Glutamax, NEAA and 55 μM 2-ME, with 10 ng/ml BDNF (R&D), 10 ng/ml GDNF (R&D). See ^15^ for full details.

### Chromatin immunoprecipitation

Native ChIP was carried out as previously described ^13^. For chromatin pull-down, the following antibodies were used: anti-H3K4me3 (Cell Signaling, Cat#9751S), anti-H3K27ac (Active Motif, Cat# 39133), anti-H3K27me3 (Cell Signaling, cat# 9733S) and H3K4me1 (Diagenode, cat#C15410037). ChIP-libraries and sequencing were performed at the Yale Center for Genome Analysis. Paired-end DNA sequencing (2×100bp) was performed on an Illumina Hi Seq2000 to an average depth of 40M reads per sample. Organoids for ChIP-seq and bulk RNA-seq were collected in parallel from the same batch of differentiation.

### RNA extraction and sequencing

Organoids samples were collected at TD0, TD30 and TD60. Total RNA extraction was performed from at least 20 dissociated organoids per sample using Arcturus PicoPure RNA isolation kit (AppliedBiosystems, cat. 12204-01). Library preparation was performed with rRNA depletion and sequenced on an Illumina Hiseq/Novaseq to obtain an average depth of 40M reads per sample (paired-end 100bp reads) at the YCGA. Organoids for bulk RNA-seq and ChIP-seq were collected in parallel from the same batch of differentiation.

### Generation of stable line expressing Zim3-dCas9

CRISPRi relies on the expression of a nuclease-deactivated dCas9 fused to a strong transcriptional repressor, the KRAB (Krüppel-associated box) domain, which recruits endogenous factors that trimethylate histone H3 on Lys9 (H3K9me3) to silence the activity of a region targeted by specific guide RNAs (gRNAs) ^29,30^. The PiggyBac transposase system was used to establish a line expressing Zim3-dCas9 for CRISPRi library screening. Vector iF53 PB-Zim3-dCas9-mScarlet-Hygro (Addgene #204726) was obtained from Michael Ward (NIH/NINDS) and used to transfect the control male iPSC line RDH913-03#9 (generated in house ^42^). 0.8 million single-cell suspension was plated in 1 well of a 6 well plate coated with Matrigel and mTeSR plus was supplemented with Y compound (5µM). After 4 hours of plating, cells were transfected by Lipofectamine Stem Transfection Reagent (STEM00008, Thermo Fisher Scientific) with total of 3ug DNA plasmid cocktail of transposase and transposon (molar ratio of 1:5). Cells were selected with 50 µg/mL hygromycin B (10687-010, Invitrogen), and mScarlet expression was monitored to ensure dCas9 expression.

### gRNA selection and library cloning

104 target loci were selected for the single-cell CRISPRi forebrain organoid validation experiment, which were composed of 94 GLEs (selected from the full set of GLE in**Table S2**), and 10 promoters. The 94 GLEs, each targeted with up to 10 gRNAs (median: 10 gRNAs/GLE; range: 6-10 gRNAs/GLE) consisted of two groups. First, 17 GLEs centrally involved in the EN regulatory hub in the ADS subnetwork (**Fig 3**) were selected. To ensure efficient targeting by 10 gRNAs, we trimmed each of the 17 GLEs (mean size: 9.6kb) to 2kb, by centering the 2kb window around the midpoint of the 100bp subregion of the GLE that has the highest number of TF binding sites as predicted by FIMO ^43^ using default setting and the HOCOMOVO v11 full human TF motif database. Second, 77 GLEs were selected based on the criterion that they harbor a schizophrenia-associated tag SNP (n=1249 SNPs, from Table S2 of Trubetskoy et al^44^. Similarly, because 38 of these 77 GLEs were longer than 2kb, we trimmed them to the 2kb subregions centered around the SNP it harbors to ensure efficient targeting. After selecting and trimming a subset of these 94 GLEs, we then evenly divided each target region to 10 bins to design one gRNA per bin, with guidescan2, requiring specificity score >0.2 and prioritized by efficiency score when multiple gRNAs passed the filter. This resulted in up to 10 gRNAs per GLE (median: 10 gRNAs/GLE; range: 6-10 gRNAs/GLE). The 10 gene promoter targets also consisted of two groups. First, six TFs involved in neuronal maturation or differentiation (BHLHE22, FOXG1, EOMES, NEUROD2, LHX6, LXM1B) were targeted with 10 gRNAs each from Horlbeck et al.’s hCRISPRi-v2 library^32^. Among these, LHX6 has two promoters targeted by Horlbeck et al., so we selected the top 5 ranking gRNAs per promoter. Second, four positive control genes (GRN, CDH2, TFRC, UBQLN2) were targeted with 3 gRNAs each from Horlbeck et al.’s hCRISPRi-v2 library, which serve as technical controls to evaluate the efficacy of CRISPRi repression, as they have been previously shown to be efficiently repressed by CRISPRi in iPSC-derived neurons ^33^. Finally, 100 non-targeting gRNAs (9.3% of final library) were added as negative controls from a curated set from Yao et al. ^45^. In total, the final gRNA library consisted of 1079 gRNAs.

To clone the gRNAs into a lentivirus vector, we first constructed a lentiviral gRNA expression plasmid by combining a U6-gRNA cassette containing the gRNA-(F+E)-combined scaffold sequence with an EGFP-P2A-PAC cassette into a lentiviral expression backbone (modifier version of Addgene#83925) using Gibson assembly. Next, we ordered the 1079-gRNA oligonucleotide library through Twist Bioscience in the following sequence format:

ATATATCTT GTGGAAAGG ACGAAACAC CG[19/20-bpprotospacer]GT TTAAGAGCTA TGC TGG AAACA GCATAG

And the library was cloned into the Esp3I-digested vector backbone by Gibson Assembly. The library was sequencing-verified by Next Generation Sequencing.

### Lentiviral production

7 million 293Ts were plated in the afternoon into 12 mL complete Opti-MEM* in a 10cm cell culture dish. The next morning, Lipofectamine 3000 reagent was prepared with Opti-MEM and mixed well. A DNA mix of psPAX2 (9.75ug), pMD2.G (3.25ug) and gRNA plasmid pool (4.3ug) was prepared in Opti-MEM followed by the addition of P3000. Sample was vortexed briefly. The Lipofectamine 3000 mix and DNA mix were combined 1:1 and incubated for 15 minutes at room temperature. Half the media (6 mL) was removed from the dish before adding 3 mL of L3000 / DNA mix dropwise. The cells were incubated for 6 hours before discarding media and adding 12mls of complete Opti-MEM. The supernatant was collected 24 hours after transfection, stored at 4C, and replaced with 12mls complete Opti-MEM. The supernatant was collected again, 48 hours after transfection and pooled with the first harvest. The supernatant was passed through a 0.45um filter before adding 4x LentiX concentrator. Filtered supernatant was stored overnight at 4C. The following morning, sample was spun down at 1700g for 45 minutes, aspirated and resuspended in 1/50 of the original volume of complete Opti-MEM.

### CRISPRi library infection

Four million cells of the RDH913-03#9 line stably expressing Zim3-dCas9 were transduced with the gRNA library with MOI of 5. After 24h of infection, cells were allowed to grow in virus free media. Cells were then selected with 1 µg/ml puromycin (GIBCO/Thermo Fisher Scientific; Cat. No. A1113803) for 6 days followed by 1 week in drug free media to form colonies. The line was differentiated to forebrain organoids as described previously.

### 10x genomics scRNA-seq

iPSC line infected with CRISPRi gRNA library, and organoids at TD0 and TD30 were harvested for scRNA-seq. Single-cell suspension was prepared using Accutase. Cells were processed using 10× Genomics single-cell library preparation protocol with 20,000 cells targeted per libraryusing a 10× Chromium high-throughput chip (Next GEM X) and Chromium Next GEM Single Cell 5’V3 Reagent Kits with Feature Barcode technology for CRISPR Screening (10× Genomics, Inc, Document number CG000418, Rev C). For correct cell recovery, one library was collected at undifferentiated iPSC and TD0 stage and 6 libraries were collected at TD30 stage. Gene expression libraries were sequenced at 500 M reads/library and gRNA amplified libraries at 50 M reads/library on Illumina HiSeq.

### FOXG1 LOF line

Loss-of-function (LOF) mutations in the FOXG1 gene were obtained by CRISPR-Cas9 genome editing as previously described ^46^. Briefly, a guide RNAs (gRNAs) targeting the beginning of the first and only exon of the FOXG1 genes (5’ CAGGCTGTTGATGCTGAACG3’) followed by the PAM seq 5’ AGG 3’ was used to generate FOXG1 LOF iPSC lines. Cas9/gRNA mediated indels were generated in a control iPSC male line (1123-01#3) by non-homologous end joining (NHEJ), which resulted in frameshifts and premature stop codons, thus generating FOXG1 truncated proteins and presumably FOXG1 LOF alleles. Single cell-derived edited clonal lines, each containing a homozygous or heterozygous genetic mutation in the FOXG1 locus, were generated from the 1123-01#3 iPSC edited line by the limiting dilution process and screened by Sanger sequencing to identify and validate the FOXG1 mutation in each clone. Genomic DNA from single clones was extracted, PCR was performed to amplify the targeted region, and the PCR products were analyzed by Sanger-sequencing. Upon genotyping by Sanger-seq, one of the edited clones, was also analyzed by whole genome sequencing at 30x sequencing coverage ^46^. Two homozygous mutations generated in different iPSC clonal lines were used in this work: the deletion of 10 base pairs (bps) or the insertion of 1 bp (**Fig. 6C**), both resulting in frameshift and premature stop codon, generating a 188 or 120 amino acid-truncated protein, retaining only 17 or 18 amino acids (AA) of the normal FOXG1 protein, respectively. Both versions of the truncated FOXG1 protein are depleted of all the FOXG1-functional domains, including the forkhead DNA binding domain (see schematic of the resulting FOXG1 truncated proteins in **Fig. 6C**). These FOXG1 -/- lines were compared to control FOXG1 +/+ lines, including both the founder iPSC line before editing and clonal iPSC lines where the CRISPR/Cas9-induced DNA breaks didn’t result in detectable genome edits.

### BIOINFORMATIC AND STATISTICAL DATA ANALYSIS

#### Enhancer discovery

ChIP-seq reads were mapped to the UCSC hg38 reference genome using bowtie2 ^47^. ChromHMM version 1.22 ^17^ was used to segment the genome based on the ChIP-seq profile. ChromHMM fits an emission profile for each experimental group, which we defined as organoid TD + donor ASD diagnosis. We manually determined that a Hidden Markov Model with nine hidden states provides the best fit for the data, based on how many states aligned with the roadmap epigenome emissions signature. These states were grouped into broad regulatory categories: states: 1, 8, and 9 were defined as promoter states (“Prom”), 2 and 3 as repressed states (“Rep”), and the remainder as enhancer states (“Enh”) (**Fig. 1c**).

To construct a set of candidate regulatory regions that are consistent across all experimental conditions, we adapted our previous strategy ^13^ to merge and filter H3K27Ac ChIP-seq peaks. MACS2 ^48^ was used to call “original peaks” (OP) for each ChIP-seq sample against the corresponding DNA input collected during the ChIP experiment. Peaks from different samples with at least 1 bp overlap were merged using bedtools to generate H3K27Ac consensus peaks. Consensus peaks were then annotated based on unambiguous overlap with the hidden states inferred by ChromHMM. Briefly, if at least 60% of positions in a consensus peak overlapped with a particular broad category in a given stage, the consensus peak was annotated with that category at that stage; otherwise, it was annotated as “Mix”. Stagewise annotations were consolidated into a global annotation for each consensus peak according to the following strategy: If the consensus peak was annotated as “Mix” in any stage, its global annotation was also “Mix”. Consensus peaks that were annotated as “Prom” in one stage and “Enh” in another were also annotated as “Mix” globally. Otherwise, the global annotation took the most active annotation from any one stage, i.e. “Prom” or “Enh”. “Rep” was attributed a consensus annotation only if all stages were annotated with “Rep”. To increase confidence, we further required putative enhancers (i.e. consensus peaks annotated as “Enh”) to be supported by at least four samples, in other words, each corresponding consensus peak was merged by OPs from at least four samples.

#### Linking enhancers to genes and TF

The process of assigning putative enhancers to genes involves two strategies as previously described ^13^. Briefly, transcription start sites (TSS) for all protein-coding and lncRNA genes were defined from GENCODE (v33) ^49^. Enhancers within 21kb of any TSS on the linear genome were linked to the respective genes. These were supplemented with high-confidence long-range interactions between enhancers and a 1kb region around the TSS derived from Hi-C contact matrices in human fetal cortex ^18^ and hSC lines ^19,20^ with 10-kb resolution.

For transcription factor binding site analysis, homer2 ^50^ was used to test for enrichment of 1,201 human transcription factor position weight matrices (PWMs) from JASPAR2020 ^51^ among the DNA sequence of the GLEs. Transcription factor binding sites (TFBS) with an FDR-corrected enrichment p-value less than 0.01 were considered significant.

#### Bulk RNA-seq analysis and differential gene expression

RNA-seq reads were aligned to the hg38 genome using STAR (v2.7) ^52^ and annotations from GENCODE (v33)^49^. Gene-level counts were estimated using *featureCounts* function from Subread (v2)^53^. Sequencing batch effects were removed using ComBat-seq ^54^. Gene filtering (function *filterByExpr*), normalized RPKM expression values and all described differential expressed gene tests (DEGs) were generated following edgeR pipeline (v3.26)^55^. For the different DEGs presented in the study (results in **Data S4**), log-2 fold change (log2FC) and FDR-corrected p-values were computed from filtered counts with the *glmFit/glmLRT* functions, with “organoid stage” (TD0/30/60) and “family” used as covariates when appropriate (see **Data S1 for samples metadata**).

#### Differential enhancer activity

Similar to RNA reads, H3K27ac read counts in all gene linked enhancers (173,400 GLEs) estimated using featureCounts v2.0.3 ^53^ were considered a measure of enhancer activity and tested for differential activity using edgeR (v3.4). For any given contrast between groups, enhancers with low activity were filtered out using the *filterByExpr* function in R and differential activity was evaluated using *glmFit* and *glmLRT* functions in edgeR.

#### Enhancer-gene regulatory network

For each downstream gene target of a GLE, a pairwise Spearman correlation was computed between the GLE activity (H3K27ac ChIP-seq counts normalized as RPKM using EdgeR) and the expression of the target gene (RNA-seq RPKM). P-values were adjusted using the Benjamini-Hochberg method. Similarly, regulatory impact of TF on each GLE presenting the cognate TFBS was estimated as the Spearman correlation between expression of the TF transcript (RNA-seq RPKM) and the activity of the bound GLE (H3K27ac ChIP-seq RPKM). For both GLE-to-gene and TF-to-GLE, putative regulatory relationships were retained if passing FDR < 0.01 and absolute correlation coefficient > 0.25. The selected relationships were assembled into an *enhancer-gene regulatory network* (directed graph, available in **Data S2, T2-3**). Visualization and network-based computation were done using the R packages igraph (v1.4.3), tidygraph (v1.2.3) and ggraph (v2.1).

#### Synthetic TF-gene-regulatory network

The synthetic TF-to-gene regulatory network was built from the original enhancer-gene regulatory network so that TF are connected to genes through at least one identified enhancer element presenting the cognate TFBS. First, cases where both the TF-GLE and the GLE-gene edges existed (so both having significant correlations) were retained to link each TF to each of its downstream genes (see **Fig. S2** for the number of cases of regulatory cascades considered and retained). Then, Spearman’s correlation between the TF and its downstream gene(s) expressions (both RNA RPKM) were calculated and retained if FDR < 0.01 and absolute correlation coefficient (r > 0.25 or r < −0.25). Edges were then retained if the sign of the TF-to-gene correlation was coherent with the TF-to-GLE and GLE-to-gene correlation signs, which is equivalent to:

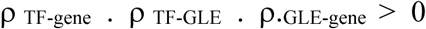

For plotting, the correlation between the genes was used to compute a positive edge weight in the final gene regulatory network with weight = r_TF-gene_ /2 +0.5 (or its inverse depending on the algorithm weight definition). Networks plotted in the different figures were obtained using either Sugiyama layout, which limits the number of edge crossing in the final hierarchical plot or Kamada-Kawai layout in igraph/ggraph, which effectively repulse strongly in the final plot genes connected by a negative correlation edge (repressive) than genes connected by a positive correlation.

#### TF prioritization by TF perturbation score

For integrating gene regulatory network with results obtained from differential gene expression results (DEG, by edgeR), we were inspired by the *AUCell GRN* method for scRNA-seq data (part of the SCENIC suite ^56^) which evaluates the activity of the TF based on the downstream targets expression enrichment in each cells. To infer similar effect using bulk data and DEG results, we used an adaptation of the *geometric perturbation index*^57^. We exemplified the use of this **TF perturbation score** in **Fig. S6a-b**. For a TF with N targets in the GRN, its perturbation score for a given DEG result was defined as:

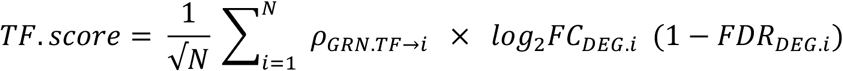

where for the i^th^ target, r_GRN.TF->i_ is the Spearman’s correlation coefficient between the TF and the i^th^ target expression level across samples as described above (corresponding to the edge’s weight in our GRN, with: −1 < r < 1). The DEG’s log_2_FC value of the downstream gene is weighted by its false non-discovery rate (1-FDR_DEG_) to downplay the effect of genes with low significant change without having to set a threshold on the significance. For each edge, the product of the sign of the logFC (−1 for DOWNregulation and +1 for UPregulation) with the sign of the correlation coefficient (+1 for activation, −1 for repression) effectively evaluate if the direction of change is compatible with the action of the TF without having to treat separately activation/repression and upregulation/downregulation (see **Fig. S6b** for examples). This generates a score with biological interpretation: a positive TF perturbation score indicates an upregulation of the TF actions described in the GRN (i.e. TF activating/repressing actions on its downstream genes), a near-zero score suggest little implication of the TF in the observed DEG results, and a negative score indicates a downregulation of the TF action. The score only considers the differential expression of the downstream targets and not of the TF itself and use the built GRN to infer the degree to which a change in the TF activity could have explained those observed differential expression.

This score was used in **Fig. 2B** with timeDEG (using in that case only activating edges with r >0 to compute the score to facilitate interpretation, see **Fig. S6C** showing all result); **Fig. 2E** with celltype-DEG (see below); **Fig. 3B** with macro-ASD DEG; **Fig. 3D** with normo-DEG; and **Fig. 6G** with FOXG1 loss of function DEG. For plots, we compared the TF perturbation score with the differential expression of the TF itself (dot blue-red color gradient), the number of target N (dot size) and the fraction of those downstream targets that meet strict DEG criteria (regardless of direction of change, cutoff: FDR < 0.05 or 0.01 and absolute log_2_FC > 0.25). Note that using the square root of N stems from the original geometrical definition of the metric and has here for effect to give higher score to genes with larger number of targets. For the specific case of **Fig. 2e**, we applied the TF perturbation score using cell type-specific DEGs computed in our scRNA-seq dataset. For cell type-specific DEGs (**Data S5**), the logFC and BH-adjuster p value were computed in Seurat (v3) using the *FindMarkers* function which performs differential expression test by comparing cells from each cell type to the remaining cells using a Wilcoxon’s rank sum test.

#### Intersection of enhancers regions external datasets

Overlap of enhancer consensus peaks identified in our data with gene enhancer external datasets were evaluated using BedTools (*intersectBed*) ^58^. From this present study we used 3 different level: the full 367,587 consensus peaks annotated as “active enhancer” in any of our sample group (“all enhancers”), the 173,347 gene-linked enhancer subset (GLE, following the gene linkage step, **Data S2,T1**) and the 42,035 GLEs that show significant correlation with any linked gene or with any TF with TFBS as described above, **Data S2,T2-3**) For Screen enhancer regions from the ENCODE project ^22^, “candidate enhancer” coordinates in hg38 was downloaded from https://screen.wenglab.org/ (v3). For Genehancer dataset ^23^, the GeneHancer_v5.14 was obtained through the Genecard website. Enhancer dataset from Amiri et al. previously generated from fetal brain, iPSCs and organoids at TD0, TD11,TD30 by our lab was also used ^13^.

Intersection with ATAC-seq/scATAC-seq external datasets were performed using BedTools ^58^ considering a minimum of 20% overlap between the length of our enhancers and the length of peaks from the external dataset. For **Fig. 1**, the resulting data was plotted using the ‘UpSetR’ package in R ^59^.

#### Intersection with Single cell ATAC-seq peak from parallel organoid data

To validate the enhancer activity in single cells, we intersected our list of enhancers with peaks called from 29 scATACseq samples using BedTools ^58^ (**Data S5**, T2). To process scATACseq data, Cell Ranger ATAC 2.0.0 ^60^ along with pre-built reference arc-GRCh38-2020-A-2.0.0 was applied to each sample from calling cells to generating a peak-cell count matrix. Signac vignette (https://stuartlab.org/signac/articles/pbmc_vignette.html) ^61^ was followed to filter the count matrix of each sample with the following cut-offs: cells containing at least 200 features, features present in at least 10 cells, log10 of fragments in peak regions per cell within two standard deviation of the mean, nucleosome signals between 2 and 4, TSS enrichment greater than 2, reads in peaks over 15% and less than 5% of peaks overlapping with black list. scATACseq data were annotated using label transfer method implemented in Seurat ^62^ along with our previously annotated scRNAseq datasets as reference ^15^. Next peaks per library or per cell type were called using MACS2 as recommended by Seurat developers.

#### TF interactome

TF-TF interactome graph based on shared gene targets between TFs in the GRN (**Data S3**). Specificity of the overlap was assessed by a connection specificity index (CSI) which is independent of the direction of the TF action (i.e. repress/activate) but evaluates if the overlap in regulon is higher for that pair of TF than with all other TFs (CSI) ^63^. Collaboration vs competition was assessed by the Pearson’s correlation coefficient (Inter.cor) between the TF’s weights across all gene targets (negative values indicate opposite actions on the downstream genes on average).

#### scCRISPRi data analysis

CRISPRi gene expression and gRNAs libraries were processed with Cellranger (version 9.010X Genomics) A threshold of 5 gRNA UMIs/cell (--min-crispr-umi) was used to assign gRNAs per cell during Cellranger count function. For the TD30 dataset, all libraries were aggregated before proceeding with data analysis. Cellranger files were further processed with Seurat (V 5.0.1) for filtering low quality cells, data normalization, and UMAP projection. To test genes susceptible to be affected by each gRNA in *cis*, chromosome position for all gRNAs in hg38 version were extracted using *BSgenome* package (v1.70.2) and genes located in a 1Mb upstream and downstream window around each gRNA were fetched using *getBM*() function of *biomaRt* package (v2.58.2). Distal genes linked to enhancers beyond 1Mb regions previously identified from HiC data were added to the list of tested genes for each loci. Pairing of gRNAs and unique hgnc symbols was carried out and these pairs were subjected to ‘SCEPTRE’ tool analysis ^35^. Counts matrices for genes and gRNA were extracted from Seurat object and used to construct a single-cell covariate matrix with non-zero expression in gRNA and gene expression libraries. These data sets were exported using packages, sceptre (v0.10.2) and ondisc(v1.2.0) for processing with ‘NextFlow’ based sceptre pipeline. sceptre pipeline was run with ‘high’ moi parameter, using positive control pairs as gRNA targeting the TSS of genes. Discovery pairs were selected based on genes falling in 1Mb window of respective gRNA. NTCs were tested against all genes from discovery pairs and calibration was performed. Multiple testing correction using Benjamini-Hochberg method was carried out to adjust the p-values and adj_p_value <0.1 was considered as significant. Change in cell fate was calculated by assessing percent alteration in each cell type for gRNAs obtained from sceptre analysis (p_value <0.05). Then mean percent alteration was calculated for each locus in all cell types. zscore was computed as following

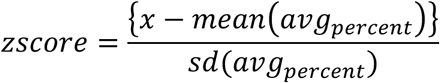

Further, p-value was calculated using a two-tailed z-test and adjusted with BH correction.

Seurat *FindMarkers*() function using Wilcoxon’s rank sum test was used to identify genome-wide DEGs for gRNAs targeting the TSS of a TF or for gRNAs targeting an enhancer with significant downregulation of a TF expression (after SCEPTRE analysis). All gRNAs targeting particular loci were combined into one group and background cells (without carrying gRNA targeting TSS of any TF) were used as control. FDR <0.05 was considered as significant alteration of gene expression.

