## Supplementary material for "Enhancer-driven gene regulatory network of forebrain human development provides insights into autism": Suppplemental Figures

Alexandre Jourdon *et al.*

**This PDF file includes:** Figs. S1 to S10

**Figure S1.** General metrics related to the linkage between genes, gene-linked enhancers (GLE) and TFs; related to Figure 1.

**Figure S2.** Inferring regulatory relationship by correlating enhancer activity and gene expression for each TF-GLE and GLE-gene linkage; related to Figure 1.

**Figure S3.** Examples of local enhancer regulatory network upstream for specific genes; related to Figure 1.

**Figure S4.** TFs' interactome delineates collaborations and competitions between TFs; related to Figure 1.

**Figure S5.** Differential expression between early (TD0) and late stages (TD30/60) confirms the progression of neurogenesis and its relationship with cell type specification; related to Figure 2.

**Figure S6.** Using *activity perturbation score* as a metric to evaluate activity changes of each TF between any two conditions based on expression changes in their regulons; related to Figure 2.

**Figure S7.** Differential gene expression between ASD probands and controls; related to Figure 3.

**Figure S8**. Differential activity of gene linked enhancer (DAE) and linked genes between ASD proband and control by head-size cohort and time point; related to Figure 3.

**Figure S9**. CRISPRi validation for predicted enhancers in forebrain organoids; related to Figure 5.

**Figure S10.** FOXG1 downstream network in forebrain organoids; related to Figure 6.

**Other supplemental information: Table S1- S5 (uploaded as separate Excel files)**

**Table S1. Sample metadata; related to Figure 1 and Figure 5**. For T2-T5, NDA locations of fastq files are listed in columns "data.file". R1 and R2 are Read 1 and Read 2 from paired-end sequencing. Multiple R1 and R2 files indicate that a library was sequenced in multiple lanes or multiple times. I1 and I2 are sample indexes.

**Table S2. Gene linked enhancers; related to Figure 1, Figure S1, Figure 2**. T1, List of all gene-linked enhancer region identified in the study. T2, List of all regulatory links connecting enhancer to genes and TFs to enhancers. T3, List of all enhancer and genes connected in T2. T2 and T3 can be used to generate the enhancer regulatory network.

**Table S3. Gene Regulatory Network. related to Figure 1**. T1, list of all edges connecting TF to their downstream genes and associated weight. T1 can be used to generate the GRN used for plots and analysis in the study.

**Table S4. Differential test results; related to Figure 2, Figure S5, Figure 6, Figure S7, S8.** T1, timeDEG.pDEG. log fold change and FDR-corrected p values for differential expression test of TD30/60 vs TD0 (timeDEG) and ASD vs Ctrl at TD0 and TD30/60 (pDEG). T2, pDAE.TD3060. log fold change and FDR-corrected p values for differential activity test of all gene linked enhancers for ASD vs Ctrl at TD30/60 (pDAE). T3, FOXG1.LOF.DEG. log fold change and FDR-corrected p values for differential expression test of FOXG1-LOF vs control lines at TD30/60 (FOXG1 LOF-DEG).

**Table S5. Cell type-specific data; related to Figure 2, Figure S5.** T1, Cell type markers from scRNA-seq data. T2, scATAC-seq peak overlap. Number of organoid scATAC-seq samples showing peak overlap with gene-linked enhancers defined by bulk ChiP-seq for all cells or by cell types. T3, scATAC-seq_libraryQC. Quality control metrics of scATAC-seq libraries used for peak overlap (from cellranger).

**Table S6. CRISPRi data; related to Fig. 5.** T1, grna id, sequence and its corresponding CONP.T2, SCEPTRE analysis results at TD30 stage. T3, Identification of enhancer-gene links within 1MB window for enhancers having at least 1 gene downregulated upon perturbation. T4, Cell fate changes due to CRISPRi mediated perturbation. T5, Effect of knockdown of TF on respective downstream targets predicted by GRN.

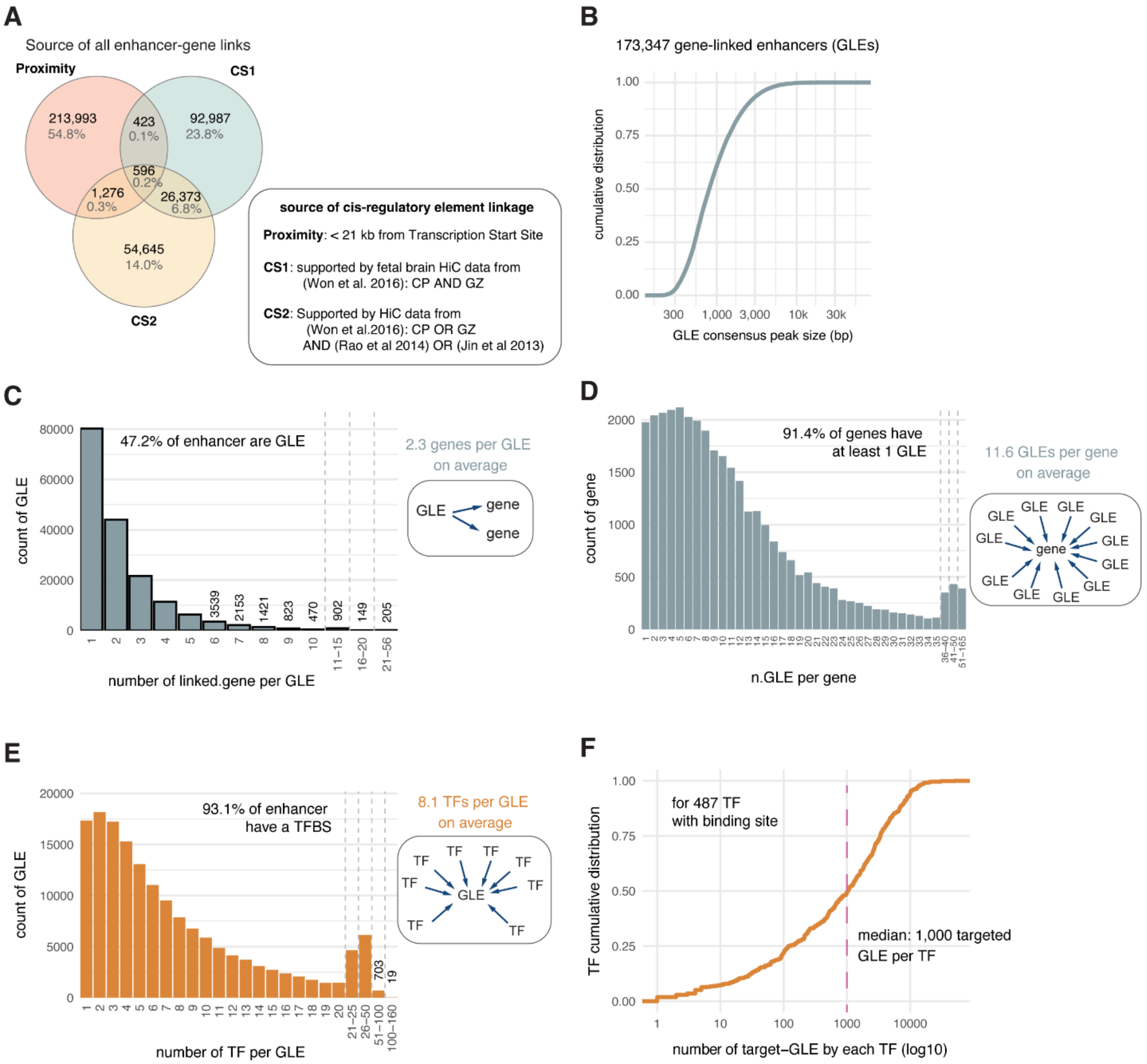


**Fig. S1. General metrics related to the linkage between genes, gene-linked enhancers (GLE) and TFs. Related to Fig. 1.**

**A.** Venn Diagram showing the sources used to link each GLE to downstream gene(s). See boxed description: CS1= confident set 1 supported by Hi-C linkages from both cortical plate (CP) and germinal zone (GZ) datasets from 3 human fetal brains (Won et al 2016, ref #9); CS2= confidence set 2, supported by CP OR GZ set (Won et al 2016, ref#9), AND by either Rao et al 2014 (ref# 11) OR Jin et al 2013 (ref # 10) human stem cell data.

**B.** Cumulative distribution of sizes of all GLE consensus peaks. Wider GLE peaks were obtained by merging sets of overlapping peaks throughout samples (see **Methods**).

**C-E.** Histograms showing the distribution of number of linked genes per GLE (C), number of linked GLE per gene (D), number of unique TF targeting each GLE (though the detection of TFBS, E).

**F.** Cumulative distribution of the number of targeted GLE per TF. Pink line=median. X-axis is in log10 scale.

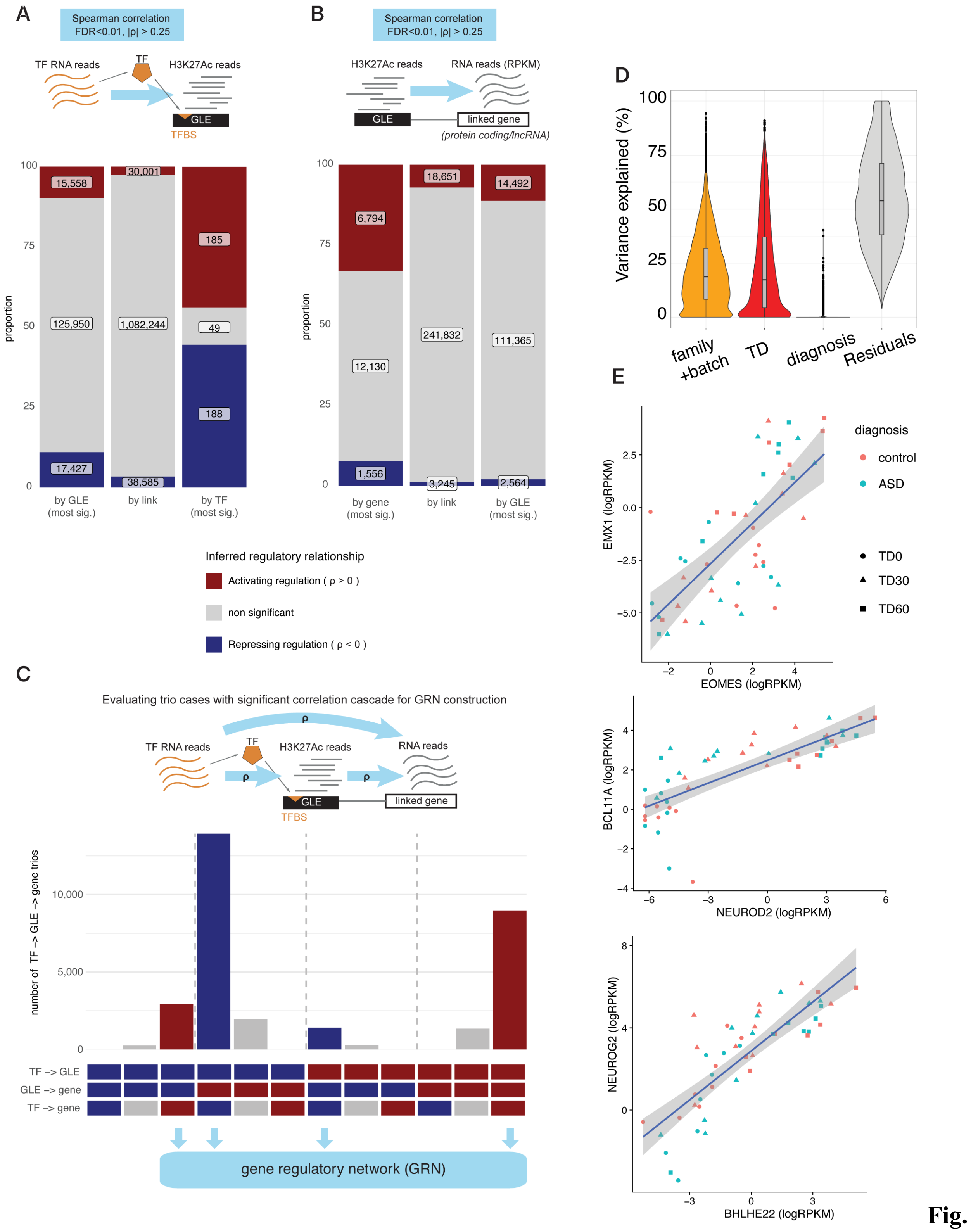


**Fig. S2. Inferring regulatory relationship by correlating enhancer activity and gene expression for each TF-GLE and GLE-gene linkage. Related to Fig. 1.**

**A.** Bar plots showing the classification of TF-to-GLE regulatory relationships as evaluated by significant correlation between GLE's H3K27ac reads level and TF's RNA reads level (two-sided

Spearman’s correlation coefficient |ρ| > 0.25 and FDR-corrected p-value < 0.01). “by GLE (most sig.)” = status of GLE according to its most significant correlation (since GLE can be connected to multiple TFs and vice-versa, only the link with the highest significance is considered here to categorize the element as activated or repressed); “by link” = status of each unique link (i.e,. TF-enhancer pair), “by TF (most sig.)” = status of each TF according to their most significant correlation.

**B.** Bar plots showing the proportion of GLE-to-gene regulatory relationships as evaluated by significant correlation between H3K27ac reads level and linked transcript RNA reads level (Spearman’s correlation coefficient |ρ| > 0.25 and FDR-corrected p-value < 0.01). “by gene (most sig.)” = status of gene according to their most significant correlation (i.e., activated or repressed); “by link” = status of each unique GLE-gene link evaluated; “by GLE (most sig.)” = status of each GLE according to their most significant correlation.

**C.** Bar plot showing the type of regulatory cascade identified for TF-GLE-target gene trios. Trios are separated by regulatory direction for each step, including the presence of an overall significant correlation between the transcript RNA level of the TF and the putative downstream gene in bottom row, as indicated by colors below the plot (red=activating, blue=repressing as in **a**, **b**). Note the absence of trios with *inconsistent* relationships. An example of inconsistent relationship would be a case where the TF is negatively correlated with the GLE and the GLE is positively correlated with the downstream gene but where the TF is positively correlated with the downstream gene suggesting an inconsistency in the cascade. For building the gene regulatory network (GRN) linking directly TF to genes, only trios with consistent correlations cascades were retained (bottom arrows), removing the cases where the TF and the downstream genes expression levels were not correlated with each other.

**D.** VariancePartition analysis on RNA expression indicated that the larger fraction of variance susceptible to drive correlation analysis (in **c**) were mostly driven by differences across families, differentiation batches and stage differences (TD) than differences between ASD individuals and neurotypical controls (diagnosis).

**E.** Examples of correlation between expression levels of TF (x axis) and target (y axis) across samples showing limited influence of diagnosis on overall correlation (linear regression line and se is shown for reference, but note that Spearman correlation was used in **c** to measure TF to gene association).

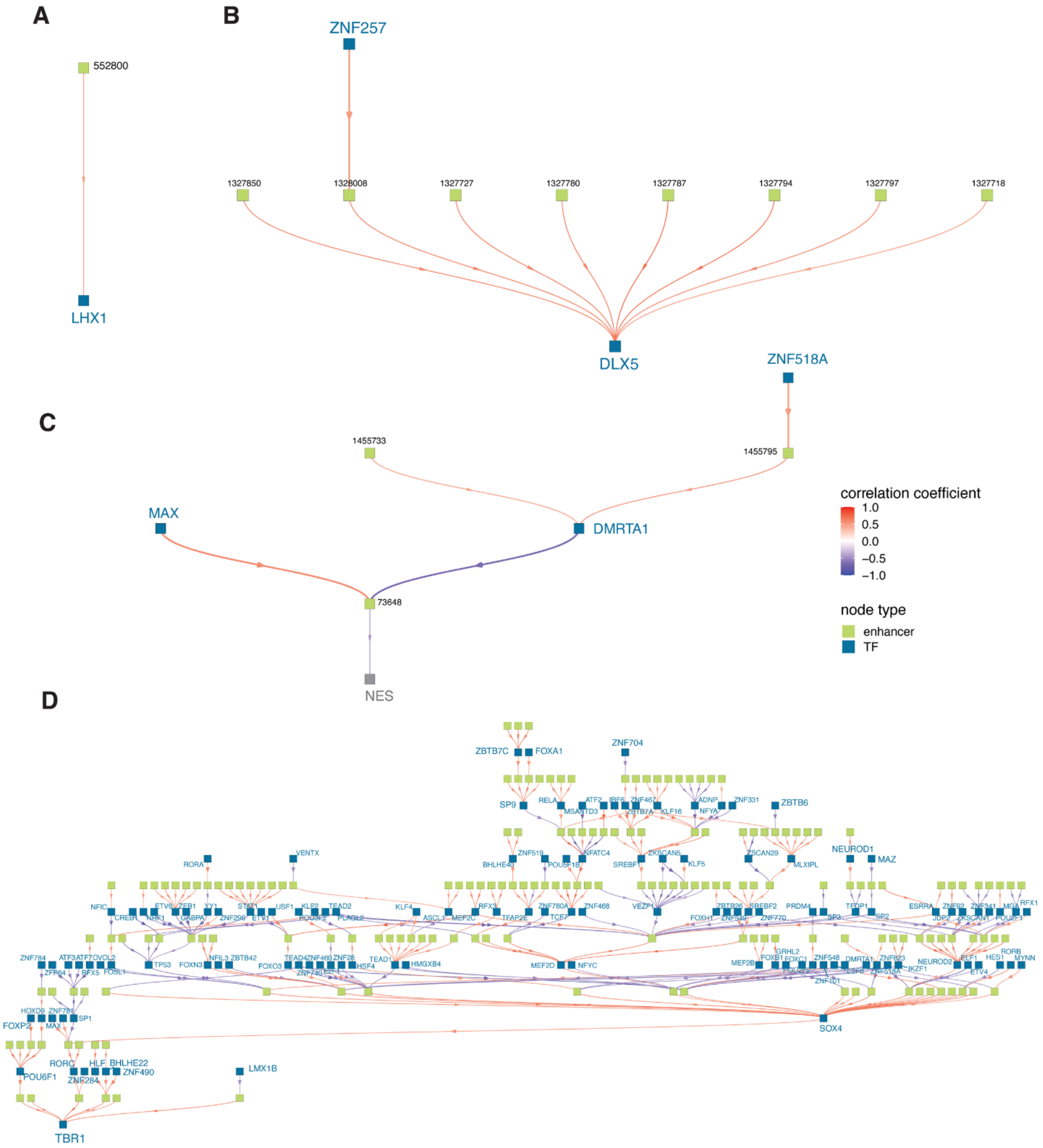


**Fig. S3. Examples of local enhancer regulatory network upstream for specific genes. Related to Fig. 1.**

**A-D.** Examples of detailed upstream enhancer-gene regulatory networks focused on *LHX1* (**A**), *DLX5* (**B**), *NES* (*NESTIN*, **C**) and TBR1(**D**). Networks **(A)-(D)** are organized by increased level of complexity. From each starting gene (bottom square), upstream cis-regulatory GLE(s) (green squares, labelled by unique ID from **Data S2**) are indicated in the 2nd layer and relationship to the target gene(s) is shown by descending arrows (activating relationships in red and repressing in blue as evaluated by correlation coefficient). When an upstream binding TF have been identified,

GLEs are then connected to upstream TFs (blue squares) identified in the network and connected by a descending edge with arrow. If upstream TFs have been themselves associated with further upstream GLEs in their own cis-regulatory region, those are plotted on an additional layer. Further TF-GLE and GLE-to-gene are then plotted in additional layers, propagating therefore the network in the upstream direction until no upstream node is identified in the network (the plot is therefore meant to be read from bottom to top, i.e., in the upstream direction). Note that in this network visualization, TF blue squares represent both the TF *gene transcript* and the TF protein as a unique element. If the gene of interest is itself a TF (e.g., TBR1, LHX1 or DLX5), its downstream network is not plotted here. For simplification of the *TBR1*-focused network, only downstream edges are displayed and arrow connecting genes upward are not displayed (i.e., putative feedback loop in the system). Note that the complexity of the *TBR1* upstream network stems in part to its relationship with SOX4 that has himself a complex upstream network.

Through this analysis of upstream nodes, the enhancer regulatory network allows to identify all TFs and non-coding enhancer regions putatively associated with the expression of a gene of interest during organoid neural development and their putative hierarchy in regulatory relationships.

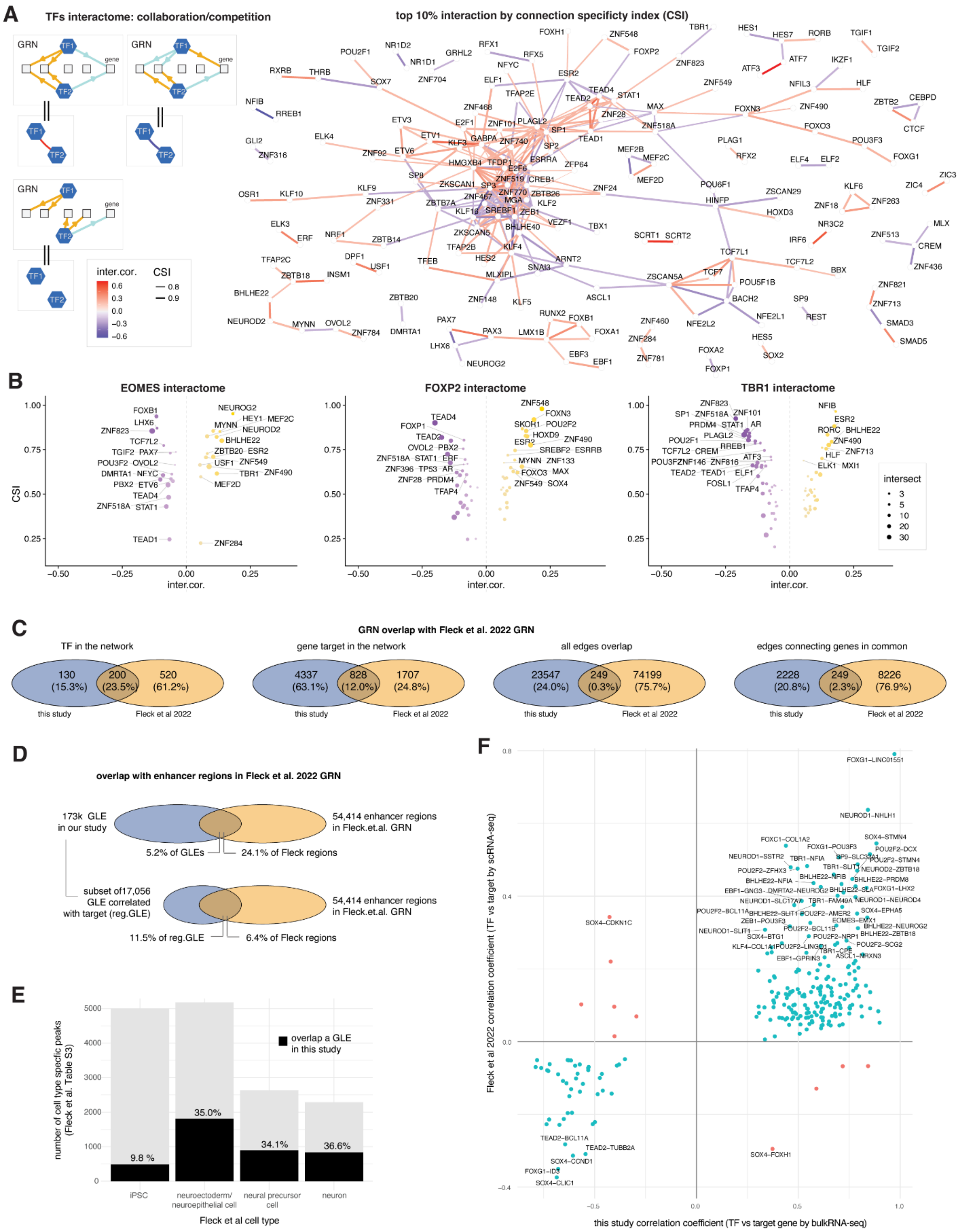


**Fig. S4. TFs' interactome delineates collaborations and competitions between TFs. Related to Fig. 1.**

**A.** TF-TF putative interactome graph based on shared gene targets between TFs in the GRN (**Data S3**). Left side schematic shows interactome evaluation to classify the TF-TF interactome (red/blue edges) based on collaborative/competitive regulation of shared targets in the GRN (yellow/cyan arrows). Specificity of the overlap assessed by Connection Specificity Index (CSI) which evaluates if the overlap in regulons is higher for that pair of TFs than with all other TFs (CSI, Bass et al 2013, ref. #41), independently of TF action (i.e. repress/activate). Collaboration vs competition was assessed by the Pearson's correlation coefficient (Inter.cor) between the TF's weights across all gene targets (negative values indicate opposite actions on the downstream genes, on average). Graph displays the top10 % specific interactions by CSI. While interactions between closely related TFs (e.g., SCRT1/SCRT2, TCF7L1/TCF7L2 pairs) could be related to a similarity in binding motif and an overlap in targeted enhancers (data not shown), interactome of TFs also included collaborators and competitors that can bind through different enhancers to influence the same genes.

**B.** Inferred interactome focused on 3 specific TFs (EOMES, FOXP2, TBR1) listing top 30 most specific collaborators (right, positive inter.cor. score as defined in (a)) and competitors (left, negative inter.cor. score) as ranked by CSI (as defined in (a)). Note that interaction can include both upstream and downstream TFs in the GRN (as interaction is evaluated by overlap in shared targets).

**C-F.** Comparison with GRN generated in Fleck et al. by scATAC-seq and scRNA-seq analysis of iPSC and early stages of organoid differentiation (< 2 months, unguided protocol). The GRN inferred using *Pando* (ref 12) was downloaded and the network were compared to the GRN inferred in the present study. Overlap in TF, gene targeted and edges (links between genes and TF) (in **c**) or in enhancer regions involved in the GRN construction (in **D**) showed a limited overlap between the 2 networks. Partial overlap can be due to different system and methods (organoid stage, single cell ATAC-seq vs bulk ChIP-seq for enhancer detection, and analytical methods). Part of the difference in enhancers included in the network stems from the inclusion of stem cells in Fleck et al., as less than 10% of iPSC-specific- regions (Fleck et al. Table S3) were present in our study (in **E**). However, while the resulting networks from both studies have only 249 edges in common (right Venn diagram in **C**), the large majority of those edges (239/249) show consistent measures in TF-to-gene correlation estimation (in **F**) suggesting that both networks capture the same regulatory actions but differ in the identification of relevant TF, gene and enhancers, due to stage of maturity and other factors, leading to different yet potentially complementary networks of organoid development. This preliminary comparison would have to be repeated on networks built from identical samples to evaluate how robust are the regulatory actions identified across epigenomic techniques, single-cell vs bulk, time points (stem cells, early organoid, late-stage organoids, etc.) and network building methods.

This difference stemmed in part from differences in the set of regulatory regions identified from chromatin marks ChIP signal (our study) and open chromatin regions signal (scATAC-seq in Fleck et al.) as well as the incorporation of earlier timepoints in the Fleck et al. study (e.g., undifferentiated iPSC and early organoids from day 4 to 61, while we assayed organoids at day 20, 50 and 80) (**Fig. S4D,E**). This ultimately resulted in a limited overlap of the two final GRNs (249 or 2.3% of TF-gene regulatory edges in common, **Fig. S4C**). However, the large majority of the common edges (239/249) show consistent measures in TF-to-gene correlation estimates between the two studies, suggesting that both networks capture the same regulatory interactions (**Fig. S4F**). This comparison agrees with the specificity of enhancer activity according to stage of development and cell type [24], suggesting that the two networks could be used in complement to chart a more complete view of regulatory actions during development. At the same time the low overlap may also point to a dependence of derived GRNs on the building methods, type of epigenomic measurements for identification of enhancers (single cell vs bulk, histone mark vs open chromatin, incorporation of HiC), and model used for regulatory inference, which will require application of these methodologies to identical datasets to be resolved.

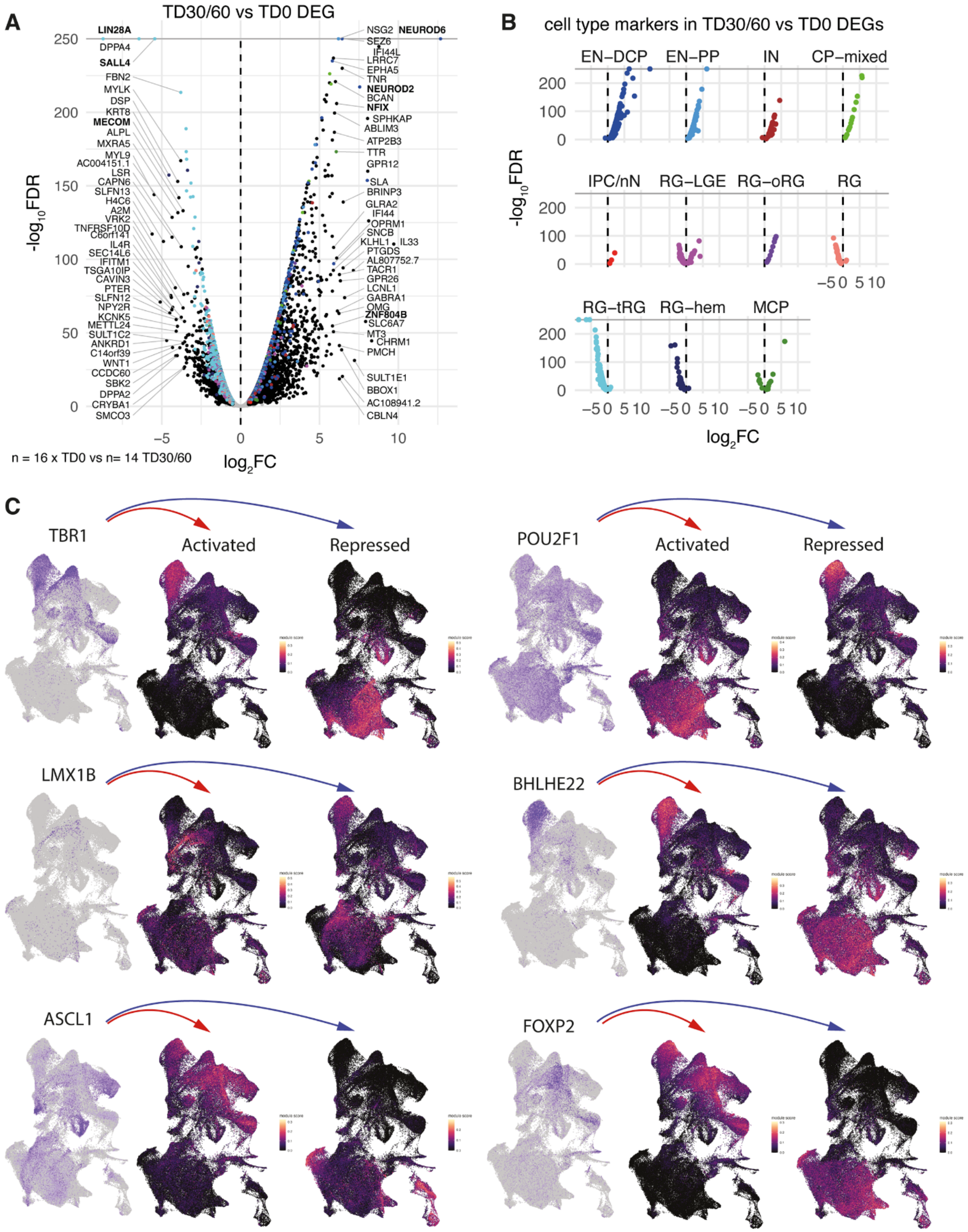


**Fig. S5. Differential expression between early (TD0) and late stages (TD30/60) confirms the progression of neurogenesis and its relationship with cell type specification. Related to Fig. 2.**

**A.** Volcano plot of the timeDEG (TD30+60 vs TD0). Dots are colored when identified as cell type-specific markers as defined by scRNA-seq analysis (cell type markers, **Data S5**). Gene names are annotated for the top 40 timeDEG as ranked by $log_2FC$ (if meeting FDR < 0.01 & $|log_2FC| > 2.5$) with TF in bold.
**B.** Isolation of cell type-specific markers from **a**. Note the different distribution of the direction of change (zero marked by vertical line) between neuronal (top) and outer radial glia (RG-oRG) cell type markers (upregulated over time) compared to early radial glia-specific markers (i.e., RG, RG-tRG, RG-hem) (downregulated over time).
**C.** Single cell expression patterns for TFs and their activated and repressed targets. Sets of 3 UMAP plots per TF showing TF expression level (left) and module scores of their downstream activated (middle) and repressed targets (right). scRNA-seq data from 70 organoid samples was used (2000 cells per library with samples from TD0, TD30 and TD60 as described in Jourdon et al. 2023) (*13*). Presented TFs had at least 15 activated/repressed targets with detectable expression level by scRNA-seq. For TF expression level, normalized expression per cell using is used (SCT -normalized counts from Seurat, blue=high and grey=low); to represent average expression for each set of targets, module score was derived (from *Seurat* R package, dark=low to purple/yellow=high). Note the broadly mutually exclusive expression patterns of repressed vs activated targets for each TF regulon.

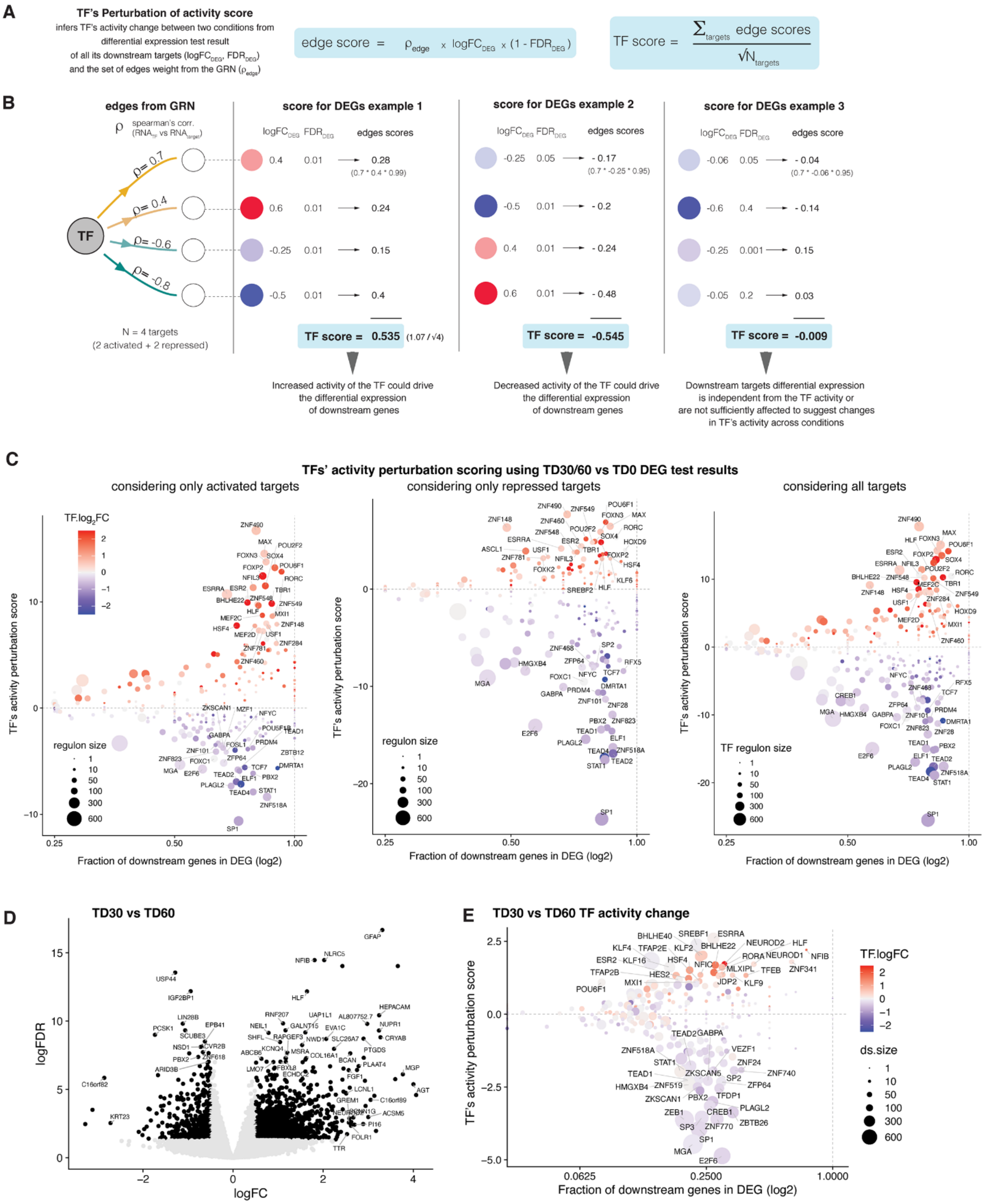


**Fig. S6. Using *activity perturbation score* as a metric to evaluate activity changes of each TF between any two conditions based on expression changes in their regulons. Related to Fig. 2.**

**A**. For each TF, an *activity perturbation score* can be computed using GRN's edges correlation values with all downstream targets included in its regulon (Spearman's correlation coefficients in our GRN) and from the result of differential expression test of the downstream targets (i.e., log2 fold change penalized by significance, as measured by the FDR-corrected p-value estimated by EdgeR differential expression test). Note that to compute the score, the DEG results do not need to be filtered for significance as significance is considered in the equation (1-FDR); the TF itself does not need to be differentially expressed and its own DEG results are not considered in the score. The perturbation score was derived from the geometric perturbation index proposed by Martin et al 2012 (*46*). See **Methods.**

**B.** Theoretical examples for 3 different sets of DEG results using a simplified TF regulon where the TF is connected to 2 activated and 2 repressed targets. Computation of the perturbation score and interpretations are shown.

**C.** For a given DEG test result such as the timeDEG (TD30/60 vs TD0), the perturbation score for each TF can be computed considering only activating (left), repressive (middle) or both types of regulatory action in their regulon (right). The perturbation score (y axis) is plotted against the fraction of the TF regulon that meet significance in the DEG results (FDR < 0.05, |log2FC| > 0.25) and the expression change of the TF itself is plotted by color.

**D,E.** A similar analysis can be performed on the TD30 to TD60 transition, which generates less pronounced changes in expression (**D**) and is driven by a different set of TF (**E**), implicating notably NEUROD2 or BHLHE22 marking ongoing excitatory neurogenesis.

Since most TF have both activating and repressive relationships they are scored in all cases and show similar ranking overall. Note that the scores here are highly concordant since the GRN was built with the same data as those used for time DEGs, but the ranking of perturbation scores extracted the most relevant TFs driving the TD0 to TD30/60 transition.

Overall, the perturbation score constitutes a versatile way to compare TF's ability to drive expression changes while exploiting both repressive and activating regulatory actions inside a complex GRN. Although there could be many different complementary or alternative metrics to derive this information, we found that the perturbation score presented here had the advantage to generate clear interpretative results. The score was used in **Fig. 2B** with timeDEG; **Fig. 2e** with cell type-DEG; **Fig. 3B** with macro-ASD DEG; **Fig. 3D** with normo-DEG; and **Fig. 6** with FOXG1 loss-of-function DEG results.

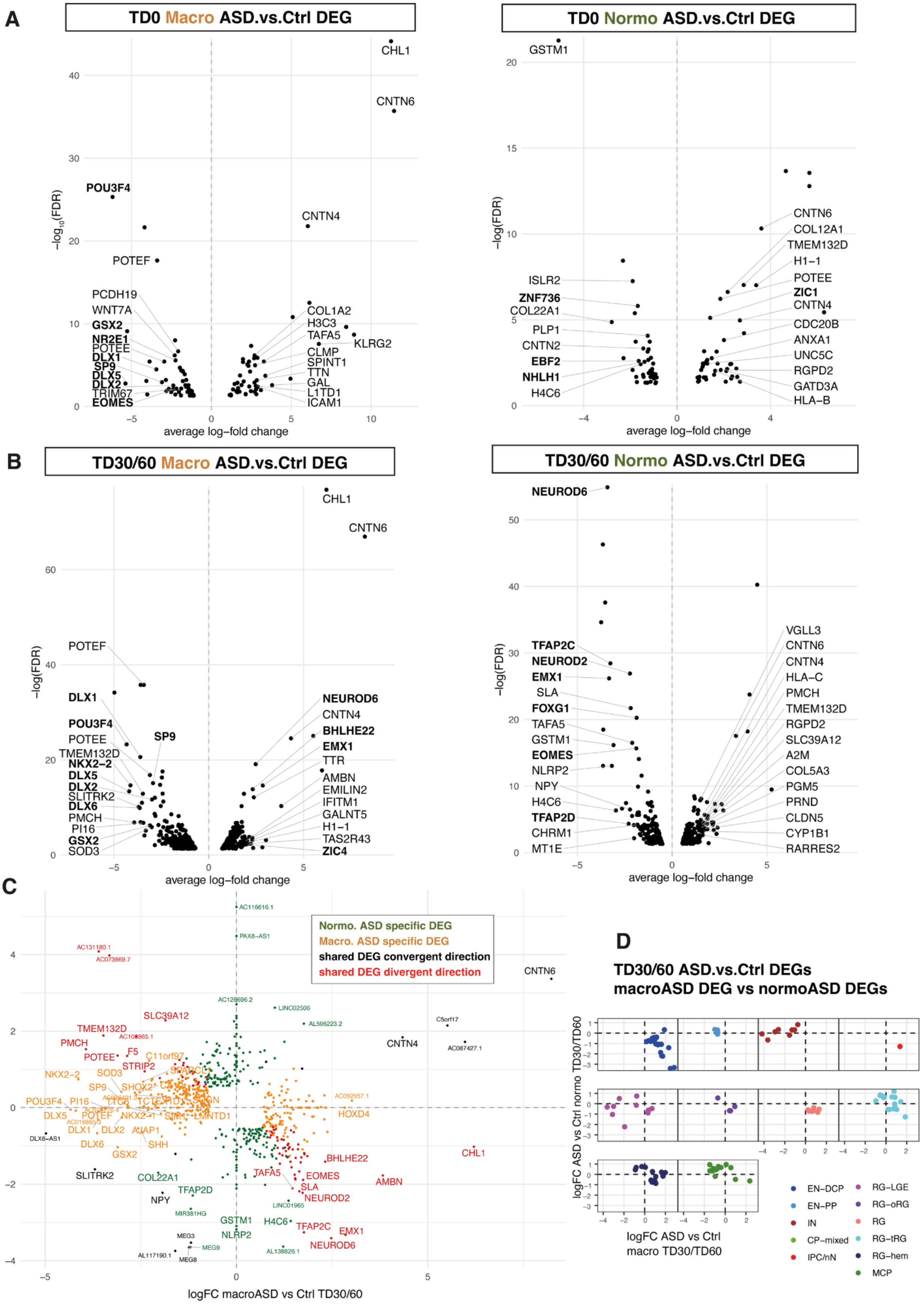
A
TD0 Macro ASD.vs.Ctrl DEG
−log10(FDR)
average log−fold change
CHL1
CNTN6
POU3F4
CNTN4
POTEF
PCDH19
WNT7A
GSX2
NR2E1
POTEE
DLX1
SP9
DLX5
DLX2
TRIM67
EOMES
COL1A2
H3C3
TAFA5
KLRG2
CLMP
SPINT1
TTN
GAL
L1TD1
ICAM1
TD0 Normo ASD.vs.Ctrl DEG
−log(FDR)
GSTM1
CNTN6
COL12A1
TMEM132D
H1−1
POTEE
ZIC1
CNTN4
CDC20B
ANXA1
UNC5C
RGPD2
GATD3A
HLA−B
ISLR2
ZNF736
COL22A1
PLP1
CNTN2
EBF2
NHLH1
H4C6
B
TD30/60 Macro ASD.vs.Ctrl DEG
−log(FDR)
CHL1
CNTN6
POTEF
DLX1
POU3F4
SP9
POTEE
TMEM132D
NKX2−2
DLX5
DLX2
SLITRK2
DLX6
PMCH
PI16
GSX2
SOD3
NEUROD6
CNTN4
BHLHE22
EMX1
TTR
AMBN
EMILIN2
IFITM1
GALNT5
H1−1
TAS2R43
ZIC4
TD30/60 Normo ASD.vs.Ctrl DEG
NEUROD6
TFAP2C
NEUROD2
EMX1
SLA
FOXG1
TAFA5
GSTM1
EOMES
NLRP2
NPY
H4C6
TFAP2D
CHRM1
MT1E
VGLL3
CNTN6
CNTN4
HLA−C
PMCH
TMEM132D
RGPD2
SLC39A12
A2M
COL5A3
PGM5
PRND
CLDN5
CYP1B1
RARRES2
C
Normo. ASD specific DEG
Macro. ASD specific DEG
shared DEG convergent direction
shared DEG divergent direction
logFC normo ASD vs Ctrl TD30/60
logFC macroASD vs Ctrl TD30/60
D
TD30/60 ASD.vs.Ctrl DEGs
macroASD DEG vs normoASD DEGs
logFC ASD vs Ctrl normo TD30/TD60
logFC ASD vs Ctrl
macro TD30/TD60
EN−DCP
EN−PP
IN
CP−mixed
IPC/nN
RG−LGE
RG−oRG
RG
RG−tRG
RG−hem
MCP

**Fig. S7. Differential gene expression between ASD probands and controls. Related to Fig. 3.**

**A-B.** Volcano plots of DEG results per cohort (Macrocephalic=macro, Normocephalic=normo) and stage (TD0 or TD30/60 = using both TD30 and TD60 samples, see **Method**). Top 15 protein coding DEGs (ranked by $log_2FC$ for each direction) and with FDR < 0.01 are annotated with TF in bold.

**C.** Confusion scatter plot of macroASD DEGs (x axis) vs normoASD DEGs (y axis) $log_2FC$ results at TD30/60 showing concordance (upper right/bottom left quadrants, black color), discordance (top left/bottom right quadrants, red color) or cohort specific (green/yellow dots) DEGs among the two cohorts. Only genes that were DEG (i.e., passing $|log_2FC| > 0.25$ & FDR < 0.05 in either cohort) were plotted. When a gene was not tested in a cohort, a dummy $log_2FC$ value of zero for that cohort was used for plotting. Gene names indicated when located at 2.5 unit from the origin (i.e. square root of ($logFC.Macro^2 + logFC.Normo^2$) > 2.5), with lncRNA in a smaller font. Color indicates the DEG category when comparing Macro and Normo results (red=shared (showing either concordant directions or discordant directions); normoASD-specific = green; macroASD specific = yellow).

**D.** Confusion scatter plots of DEG results at TD30/60 plotted as in **c** for gene with cell type-specific expression in scRNA data (i.e. cluster maker of cell types (**Data S5**)) as indicated by color. Cell type specific expression was evaluated for each of the 11 main cell types. Results show that genes expressed in EN-DCP were increased in Macro-ASD and decreased in Normo-ASD and that IN related genes were specifically downregulated in Macro-ASD.

Cell type abbreviations: EN-DCP: excitatory neuron-dorsal cortical plate; EN-PP : excitatory neuron-preplate ; IN : inhibitory neuron ; CP-mixed : mixed neurons ; IPC/nN: intermediate progenitor cell /newborn neurons ; RG-LGE: lateral ganglionic eminence progenitor ; RG: radial glia ; oRG outer radial glia ; RG-tRG: truncated RG or dividing RG ; RG-hem: cortical hem-like progenitors ; MCP: medial cortical plate cells.

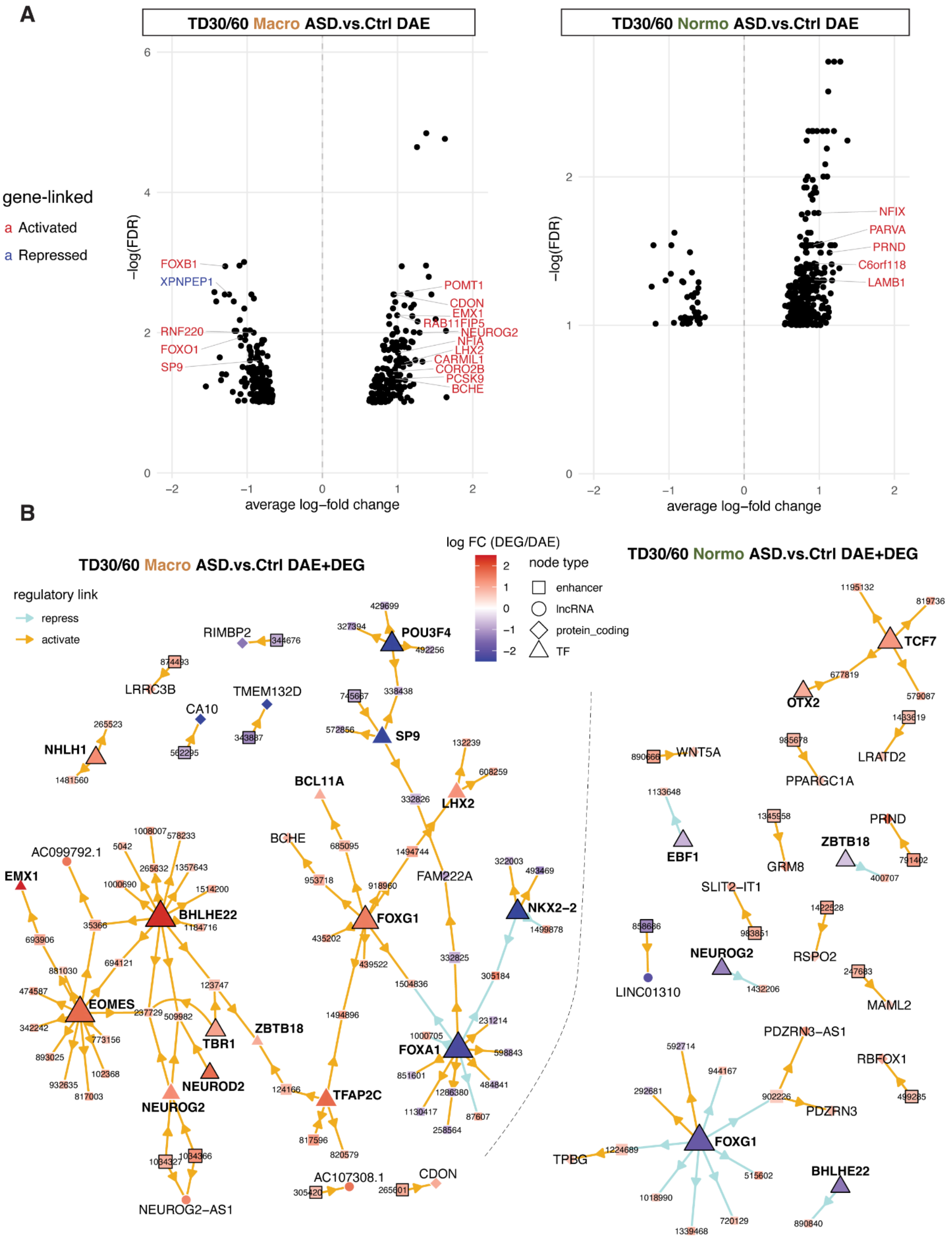


**Fig. S8. Differential activity of gene linked enhancer (DAE) and linked genes between ASD proband and control by head-size cohort and time point. Related to Fig. 3.**

**A.** Volcano plots of differentially active enhancers at TD30/60 in ASD probands vs control fathers. DAE are annotated by their linked protein coding gene when included in the regulome (i.e., with significant correlation). The relationship between the DAE with the linked gene(s) is indicated by colors: red=activating (Act.reg.) or blue=repressing (Rep.reg.) corresponding, respectively, to a positive or negative correlation between H3K27ac and transcript RNA reads across the full dataset. Note that an enhancer can be linked to multiple genes and a gene can be linked to multiple enhancers (e.g., TMEM132D). Full DAE results by gene-link enhancer ID are available in **Table S4, T2**.

**B**. DAE/DEG enhancer regulatory network. The full enhancer network was subset to plot only links connecting DAE to DEG (such link can be between a TF and a downstream GLE or between a GLE and a downstream gene target). Upstream-most elements (i.e., with no other upstream differentially active element in the network) are outlined with a black line. Note that for enhancer, a threshold of FDR < 0.1 was used.

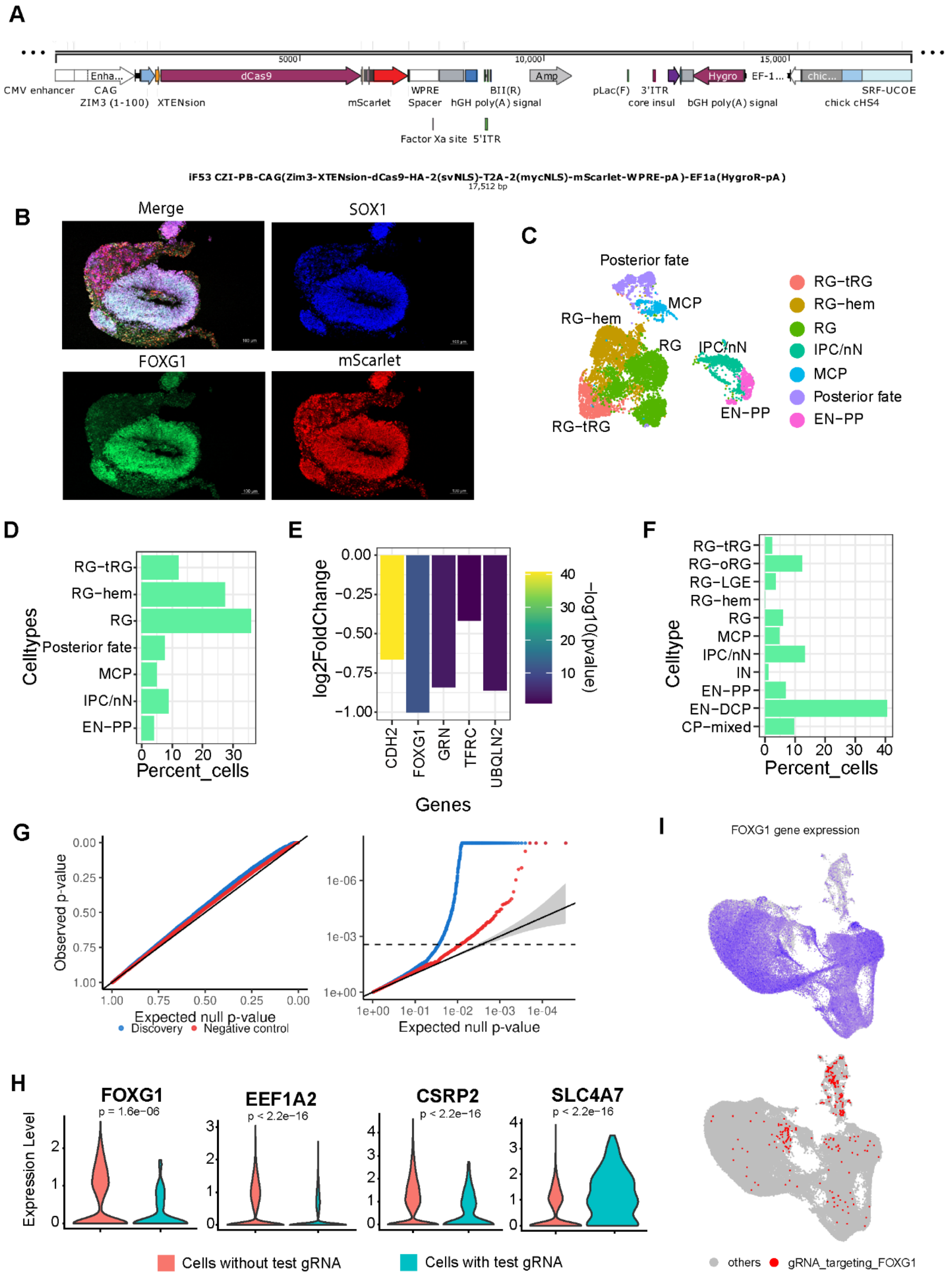


**Figure S9. CRISPRi validation for predicted enhancers in forebrain organoids. Related to Figure 5.**

**A.** Schematic of the iF53 expression vector.

**B**. Representative images of 01-01-Cas9 and RDH913-03-Cas9 differentiated into forebrain organoids, expressing dCAS9-KRAB-mScarlet, immunostained for the telencephalic gene FOXG1, and the cortical progenitor marker SOX1.

**C.** UMAP plot for TD0 stage organoid. Annotation was carried out based on cluster markers. Colour= Predicted cell types.
**D.** Bar graph representing cell type composition at TD0 stage.
**E.** Bar graph representing cell type composition at TD30 stage.
**F.** Effect of CRISPRi mediated perturbation on expression of positive control genes and FOXG1. The transcriptome of cells containing gRNAs targeting these genes were pulled for analysis and compared to transcriptome of non-targeted cells. Colour = pvalue obtained from DGE analysis.
**G**. Quantile-quantile plot for gRNA-gene pairs to test p-values pairs to non-targeting gRNAs tested against the same genes. pvalues obtained from sceptre analysis were tested here.

**H.** Examples for alteration in expression of target genes upon perturbation with gRNA. Pvalue is calculated using Wilcoxon's rank sum test.

**I.** UMAP at TD30 representing FOXG1 expression and distribution of gRNA targeting FOXG1. Red dots= cells carrying FOXG1 gRNA, grey dots= background.

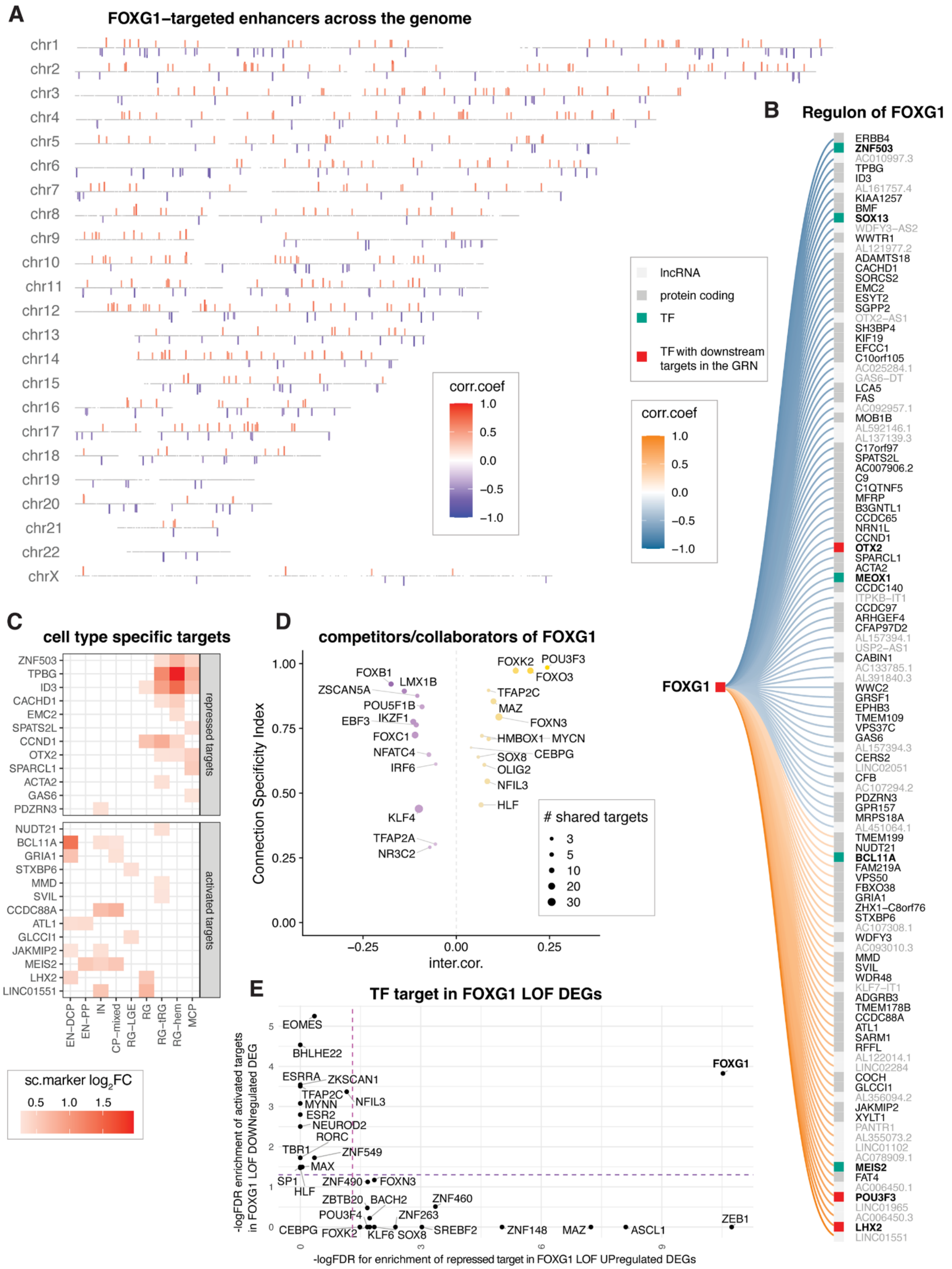


**Fig. S10. FOXG1 downstream network in forebrain organoids. Related to Fig. 6.**
**A.** Bar plot showing the genomic location of enhancers targeted by FOXG1 per chromosome, with color and bar size proportional to the correlation coefficient between the enhancer activity

(H3K27ac RPKM level) and FOXG1 transcript level (RNA RPKM) across samples. Grey trace shows the location of all gene-linked enhancer as background.

**B.** Full downstream network of FOXG1 ranked by correlation strength (edge color). The type of the downstream target is indicated by color (e.g., lncRNA, TF, etc.)

**C**. Gene targets of FOXG1 that have cell type-specific expression (marker.log2FC= DEG between the cell type and all other cells in scRNA-seq (**Data S5**), using here only positive log2FC as marker of cell type-specific expression).

**D.** Dot plot of FOXG1 interactome showing TFs that are collaborators (gold) and competitors (purple) of FOXG1 across the GRN (see Fig. S4) with the specificity of the interaction ranked by CSI (y-axis, corresponding to the fraction of all other TFs that have lesser regulon overlaps than the one observed for the evaluated pair).

**E.** Dot plot of significant enrichment of TF's targets in FOXG1 loss of function DEGs, showing both the enrichment of repressed targets in upregulated DEGs (x axis) or of activated targets in downregulated DEGs (y axis), both indicating a decrease activity of the TF following loss of function (FDR corrected p-value by one-sided Fisher's exact test).
